# LINE elements are a reservoir of regulatory potential in mammalian genomes

**DOI:** 10.1101/2020.05.31.126169

**Authors:** Maša Roller, Ericca Stamper, Diego Villar, Osagie Izuogu, Fergal Martin, Aisling Redmond, Raghavendra Ramachanderan, Louise Harewood, Duncan T. Odom, Paul Flicek

## Abstract

To investigate the mechanisms driving regulatory evolution across tissues, we experimentally mapped promoters, enhancers, and gene expression in liver, brain, muscle, and testis from ten diverse mammals. The regulatory landscape around genes included both tissue-shared and tissue-specific regulatory regions, where tissue-specific promoters and enhancers evolved most rapidly. Genomic regions switching between promoters and enhancers were more common across species, and less common across tissues within a single species. Long Interspersed Nuclear Elements (LINEs) played recurrent evolutionary roles: LINE L1s were associated with tissue-specific regulatory regions, whereas more ancient LINE L2s were associated with tissue-shared regulatory regions and with those switching between promoter and enhancer signatures across species. Our analyses of the tissue-specificity and evolutionary stability among promoters and enhancers reveal how specific LINE families have helped shape the dynamic mammalian regulome.

**HIGHLIGHTS:**

- Tissue-specific regulatory regions evolve faster than tissue-shared
- Switching promoter and enhancer regulatory roles is frequent in evolution
- LINE L1s contribute to the evolution of tissue-specific regulatory regions
- LINE L2s are associated with broad tissue activity and dynamic regulatory signatures

## INTRODUCTION

Mammalian tissues are composed of hundreds of cell types, each with its own tissue-specific gene expression program. These programs are controlled by proximal promoters and distal enhancer regions (Aguet et al., 2019).

Promoters and enhancers are traditionally considered distinct and minimally overlapping categories, although specific genomic regions can show both promoter and enhancer activity between cell types of a species (Andersson and Sandelin, 2020). Some promoters show characteristics of enhancers, such as impacting expression of distal genes (Dao et al., 2017; Leung et al., 2015); showing chromatin signatures of enhancers (Leung et al., 2015); or contacting another promoter (Jung et al., 2019). Conversely, some enhancers show characteristics of promoters by driving transcription (Andersson et al., 2014; De Santa et al., 2010; Kim et al., 2010) or functioning as alternative promoters (Kowalczyk et al., 2012). Evolutionary studies on a limited number of lineages and regulatory regions have suggested that a subset of enhancers can be repurposed to promoters across species (Carelli et al., 2018).

While gene expression divergence has been extensively characterized in mammalian tissues (Brawand et al., 2011; Cardoso-Moreira et al., 2019), the evolution of the associated regulatory regions is not well understood. Enhancer and promoter evolution has mostly been studied by comparing one mammalian tissue or cell type across several species (Danko et al., 2018; Glinsky and Barakat, 2019; Swain-Lenz et al., 2019; Villar et al., 2015). This approach is unable to compare evolutionary trends across tissues. A second approach comparing the regulatory landscapes among various tissues of mouse and human (Cheng et al., 2014; Donnard et al., 2018; Forrest et al., 2014; Vierstra et al., 2014) affords limited insights into the rate and lineage-specificity of regulatory evolution.

Nevertheless, these studies revealed that enhancers have a high rate of evolutionary turnover (Cheng et al., 2014; Danko et al., 2018; Glinsky and Barakat, 2019; Vierstra et al., 2014; Villar et al., 2015). For example, less than 5% of human embryonic stem cell enhancers are conserved in mouse (Glinsky and Barakat, 2019). Promoter regions are more evolutionarily stable (Cheng et al., 2014; Vierstra et al., 2014; Villar et al., 2015), although only around half of precise transcription start sites are conserved between mouse and human (Forrest et al., 2014; Young et al., 2015).

Tissue-specific promoter and enhancer evolution in mammals is partly shaped by transposable elements, which can contribute novel transcription factor binding sites (Bourque et al., 2018). To date, most studies have focused on the regulatory contribution of the endogenous retrovirus (ERV) superfamily of the long terminal repeat (LTR) subclass (Chuong et al., 2016; Franke et al., 2017; Jacques et al., 2013) and the short interspersed nuclear element (SINE) superfamily of the non-LTR subclass (Cao et al., 2019; Trizzino et al., 2018; Trizzino et al., 2017; Vierstra et al., 2014). Less is known about regulatory contributions of the long interspersed nuclear element (LINE) superfamily, which makes up around 20% of mammalian genomes (Platt et al., 2018). Both LINE L1s and L2s evolved before the divergence of mammals, although the L2 family is more ancient (Chalopin et al., 2015). L1s are the only elements still actively retrotransposing in mammalian genomes (Elbarbary et al. 2016). L1s are often transcribed in a cell-type specific manner (Belancio et al., 2010; Guffanti et al., 2018; Philippe et al., 2016) and there is limited evidence for their direct contribution to gene regulation (Yang et al., 1998). In human cells, LINE L2 elements are expressed as miRNAs (Petri et al., 2019) and may have regulatory activity (Cao et al., 2019; Huda et al., 2011), but it is unknown whether L2 elements play a regulatory role in other mammalian lineages.

Here, by comparing the epigenetic and transcriptional landscapes of multiple tissues and species across nearly 160 million years of mammalian evolution, we revealed new insight into the molecular mechanisms underlying tissue-specific and tissue-shared regulatory evolution. Our analyses demonstrated how promoters and enhancers can interchange regulatory signatures between species and discovered how different LINE families help shape tissue-specificity and regulatory signatures.

## RESULTS

### Mapping regulatory evolution across four tissues in ten mammals

The species selected for mapping active regulatory regions represent several mammalian clades including primates (macaque and marmoset), Glires (mouse, rat and rabbit), Laurasiatheria (pig, horse, cat and dog) and marsupials (opossum) (Table S1); all species have high quality reference genomes with extensive annotation (Yates et al., 2020).

We profiled the regulatory landscape of liver, muscle, brain, and testis - three adult somatic tissues originating from distinct developmental germ layers and one adult germline tissue. In each tissue, matched functional genomics experiments were performed in biological triplicate (with one exception, see Methods, Table S2). Chromatin immunoprecipitation followed by high-throughput DNA sequencing (ChIP-seq) was used to map three histone modifications associated with regulatory activity: histone 3 lysine 4 trimethylation (H3K4me3); histone 3 lysine 4 monomethylation (H3K4me1); and histone 3 lysine 27 acetylation (H3K27ac) (Figures 1A, 1B, S1). Libraries were sequenced to saturation to ensure reproducibility (Figure S2A, Table S3, Methods).

**Figure 1:**
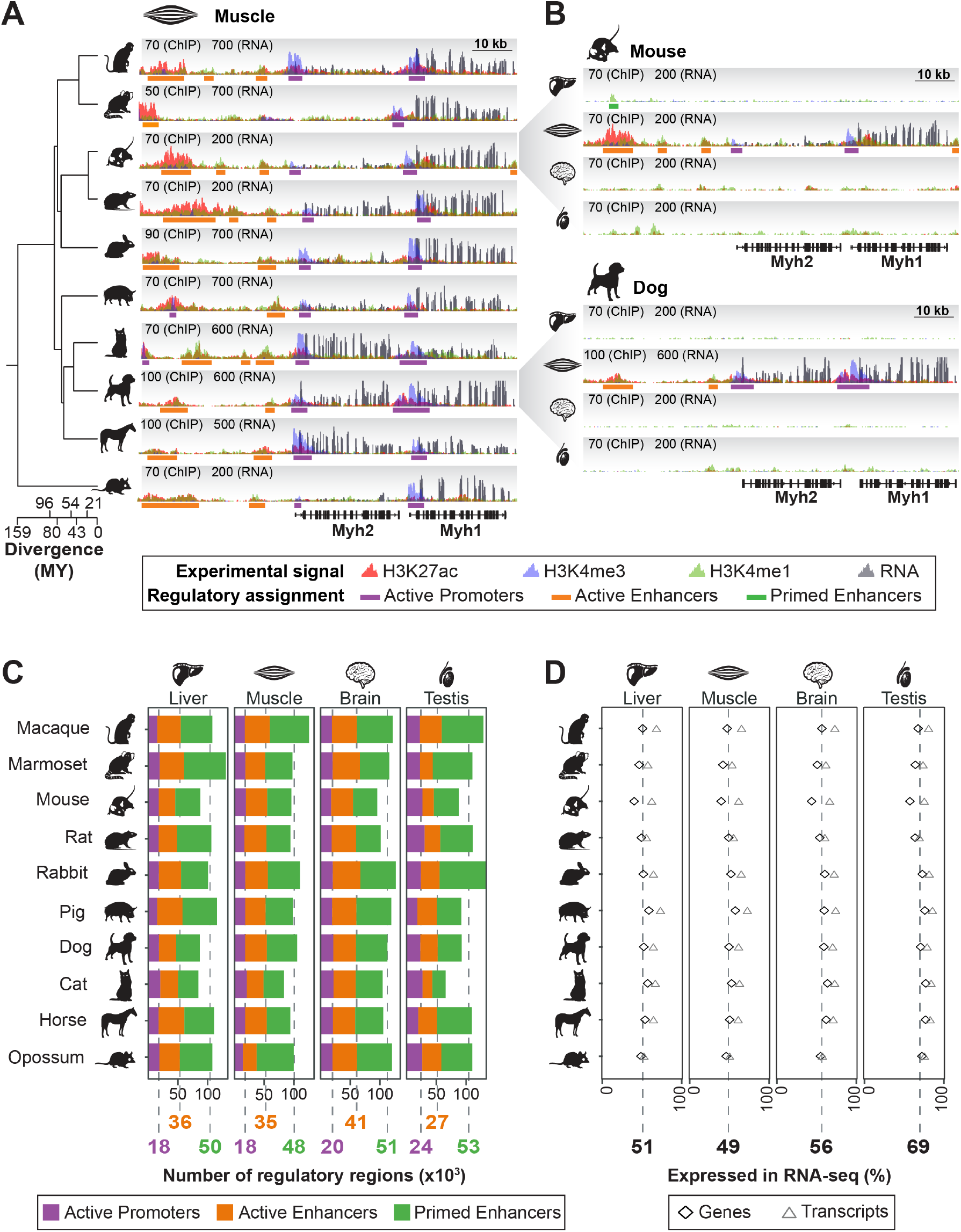
Promoter, enhancer and gene expression mapping demonstrates consistent tissue-level gene regulation in mammals. A) Example conserved, tissue-specific regulatory landscape around Myosin Heavy Chain 1 and 2 genes (*Myh1* and *Myh2*) in muscle tissue from ten mammalian species. Inset numbers are maximum read depths for ChIP-seq and RNA-seq, while phylogenetic relationships and species divergences are shown on the left. (See Figure S1 for experimental workflow.) B) For the same locus as in A), regulatory landscapes in liver, muscle, brain, and testis are shown for mouse and dog. C) For each tissue, the number of biologically reproducible regulatory regions identified is consistent across species. The average number of active promoters (purple), active enhancers (orange) and primed enhancers (green) across all species is shown below each column. (See Figure S2 for validation). D) The fraction of genes (diamonds) and transcripts (triangles) expressed in each tissue is stable across ten species. Below each column, the average percentage of expressed genes across all species is shown. Species with larger differences between the fraction of expressed genes and transcripts have more comprehensive annotation.

Active promoters were defined as regions enriched for both H3K4me3 and H3K27ac (Bernstein et al., 2005; Shen et al., 2012) (Figure 1A, 1B, S1). Active enhancers were defined as regions enriched for both H3K4me1 and H3K27ac, but not H3K4me3 (Creyghton et al., 2010; Shen et al., 2012). Primed enhancers, or intermediate enhancers, were defined as regions with reproducible peaks for H3K4me1 only (Calo and Wysocka, 2013; Schoenfelder and Fraser, 2019). These are thought to be ‘primed’ with H3K4me1 and may become readily active in response to specific stimuli (Wang et al., 2015).

To quantify genome-wide transcriptional activity, we generated matched total RNA-seq for the same samples used to map histone modifications (with rare exceptions, see Methods, Table S2). RNA-seq libraries were generally sequenced to a minimum of 20 million mapped reads for somatic tissues and 40 million for testis (Figure S1, Table S4). We used these data to improve and publicly release Ensembl genome annotations for eight species (Methods) (Aken et al., 2016; Yates et al., 2020).

From these nearly 500 matched experiments, we annotated more than 2.8 million regulatory regions in four adult tissues across ten mammalian species. This dataset captured a substantial proportion of known regulatory regions genome-wide (Figure S2B, S2C), identified thousands of novel regulatory regions (Figure S2D), and provides a comprehensive and consistent dataset for inter-tissue and inter-species analyses of regulatory evolution.

### Tissue-level regulatory and transcriptional landscapes are consistent across mammals

The number of regulatory regions identified for each tissue was largely consistent across species (Figure 1C). Liver and muscle are relatively homogeneous somatic tissues consisting mostly of hepatocytes and myocytes, respectively. Each of these two tissues expressed approximately half of all annotated genes (Figure 1D), and had on average 18,000 active promoters, 36,000 active enhancers and 49,000 primed enhancers (Figure 1C). In brain, we identified more active regulatory regions on average: 20,000 active promoters and 41,000 active enhancers. This increase is consistent with the higher gene expression we observed (56% of genes are transcribed), as well as with the greater cellular heterogeneity of brain (Darmanis et al., 2015). Indeed, the number of regulatory regions we identified in whole brain was comparable to the combined total found from profiling individual brain regions (Vermunt et al., 2016), suggesting that we effectively captured the brain regulome (Methods). Consistent with previous reports, there were twice as many active enhancers as active promoters for all three somatic tissues (Shen et al., 2012; Villar et al., 2015).

Testis is distinct from somatic tissues in that it is primarily composed of germ cells at different stages of spermatogenesis (Soumillon et al., 2013). Testis had more active promoters compared to other tissues (24,000, Figure 1C) and expressed the highest portion of annotated genes and transcripts (69%, Figure 1D), consistent with known testis transcriptome diversity (Soumillon et al., 2013; Xia et al., 2020). Testis had a lower ratio of enhancers to promoters compared to somatic tissues, suggesting a distinct regulatory landscape.

Taken together, we found that promoter and enhancer landscapes correspond to gene expression, depend on tissue identity, and are consistent across species.

### Distinctive regulatory landscapes characterize somatic tissues and testis

Within each species, we analysed the tissue-specificity (Figure 2A, Figure S3A) of enhancers, promoters, and gene expression, and then combined these into an overview for all species (Figure 2B). Consistent with previous studies (Heintzman et al., 2009; Shen et al., 2012), enhancers were mostly tissue-specific: 76% of active enhancers and 83% of primed enhancers were found in only one of the four tissues profiled. The largest group of active promoters were shared across all four tissues (37%; Figure 2B) and almost half of active promoters were tissue-specific, split between those that are testis-specific or specific to any of the three somatic tissues (25% and 23%, respectively). Transcript tissue-specificity mirrored that of active promoters. While the numbers of genes and transcripts expressed in all four tissues were similar, the number of tissue-specific expressed transcripts was 2-4 times higher than tissue-specific expressed genes, and more closely matched the number of active promoters (Figure 2B). This trend is especially evident in mouse, where the annotation is most comprehensive (Figure S3B). This suggests that tissue-specific promoters modulate alternative transcript usage.

**Figure 2:**
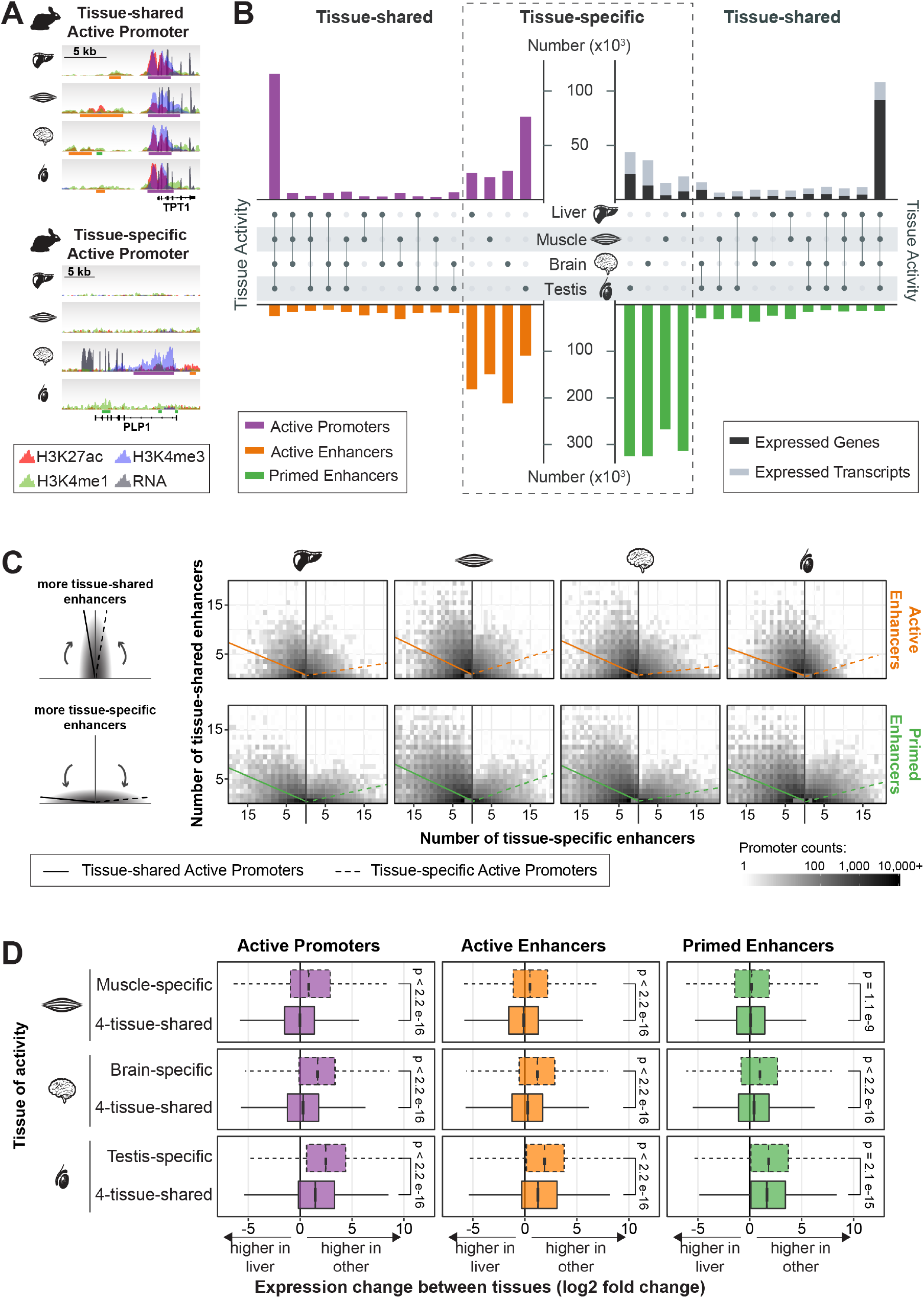
Tissue-specific enhancers are associated with tissue-specific and tissue-shared promoters. A) Example rabbit active promoters that were tissue-shared (top) and brain-specific (bottom). Y-axis scale represents read density and was normalized independently for each channel. B) Within each species, promoter activity and gene expression were distributed between tissue-specific and tissue-shared, while enhancer activity was mostly tissue-specific (see Figure S3A). The numbers are a summation across all ten study species; mouse as a representative species is shown in Figure S3B. C) Numbers of tissue-shared (y-axis) and tissue-specific (x-axis) enhancers associated with each promoter are shown for the four tissues. The schematic (right) represents two extreme examples: promoters predominantly associated with tissue-shared enhancers (top) or tissue-specific enhancers (bottom). Tissue-shared promoters (left panels) are associated with a higher ratio of tissue-shared versus tissue-specific enhancers; whereas tissue-specific promoters (right panels) are predominantly associated with tissue-specific enhancers. D) Observed expression changes between tissues for genes associated with regulatory regions in muscle, brain, and testis. The plots show the distribution of differential expression with liver as a reference (DESeq2 adjusted p-value <0.05), of genes nearest to 4-tissue-shared regulatory regions and tissue-specific regulatory regions (p-values calculated using one sided Wilcoxon test; tissue-specific expression change is greater than tissue-shared).

We investigated the association between promoters and enhancers by assigning enhancers to the nearest promoter within 1 Mb (Methods). We then examined ratios of the number of tissue-specific and tissue-shared enhancers for each active promoter. Tissue-shared active promoters typically associated with both tissue-shared enhancers and a larger number of tissue-specific enhancers (Figure 2C, left), reflecting the overall tissue-specificity of enhancers (Figure 2B). In contrast, tissue-specific active promoters associated with smaller numbers of tissue-shared enhancers and larger numbers of tissue-specific active enhancers (Figure 2C, right) compared to tissue-shared promoters. These trends were consistent for both active and primed enhancers. In testis, tissue-specific promoters associated with fewer tissue-specific active enhancers than did tissue-specific promoters in somatic tissues (Figure 2C), reflecting the overall lower number of active enhancers in testis (Figures 1C, 2B).

We assigned regulatory regions to their nearest gene and compared gene expression levels across tissues using liver as a reference (Figure 2D). Genes near tissue-shared regulatory regions showed similar expression levels across somatic tissues. In contrast, genes near muscle- and brain-specific regulatory regions had significantly higher expression in those tissues than in liver. This effect was strongest for promoters and is also evident for enhancers (Figure 2D). Genes associated with testis regulatory regions showed more pronounced expression change compared to liver, more so than the comparisons with somatic tissues, even for genes associated with tissue-shared regulatory regions (Figure 2D).

These results demonstrate that regulatory landscapes differ between somatic tissues and testis, and that tissue-specific promoters are closely associated with tissue-specific gene expression.

### Tissue-shared promoters and enhancers display enhanced evolutionary stability

The association between tissue specificity and evolutionary stability of enhancers and promoters has remained largely unexplored. Previous work in single tissues demonstrated that few enhancers are conserved across mammals (Shen et al., 2012; Villar et al., 2015), and those conserved are more likely to be active in multiple cellular contexts (Fish et al., 2017). Here, we exploited matched enhancer and promoter landscapes to identify evolutionarily maintained and recently-evolved regulatory regions (Figure S3A, Methods). The majority of tissue-shared regulatory regions (76%) were maintained in evolution, although there were also many tissue-shared regions that were recently-evolved. In contrast, most tissue-specific regulatory regions were recently-evolved (89%; Figure 3A). Across all ten species, we found 1.6 million recently-evolved regulatory regions and 1.2 million maintained regulatory regions.

**Figure 3:**
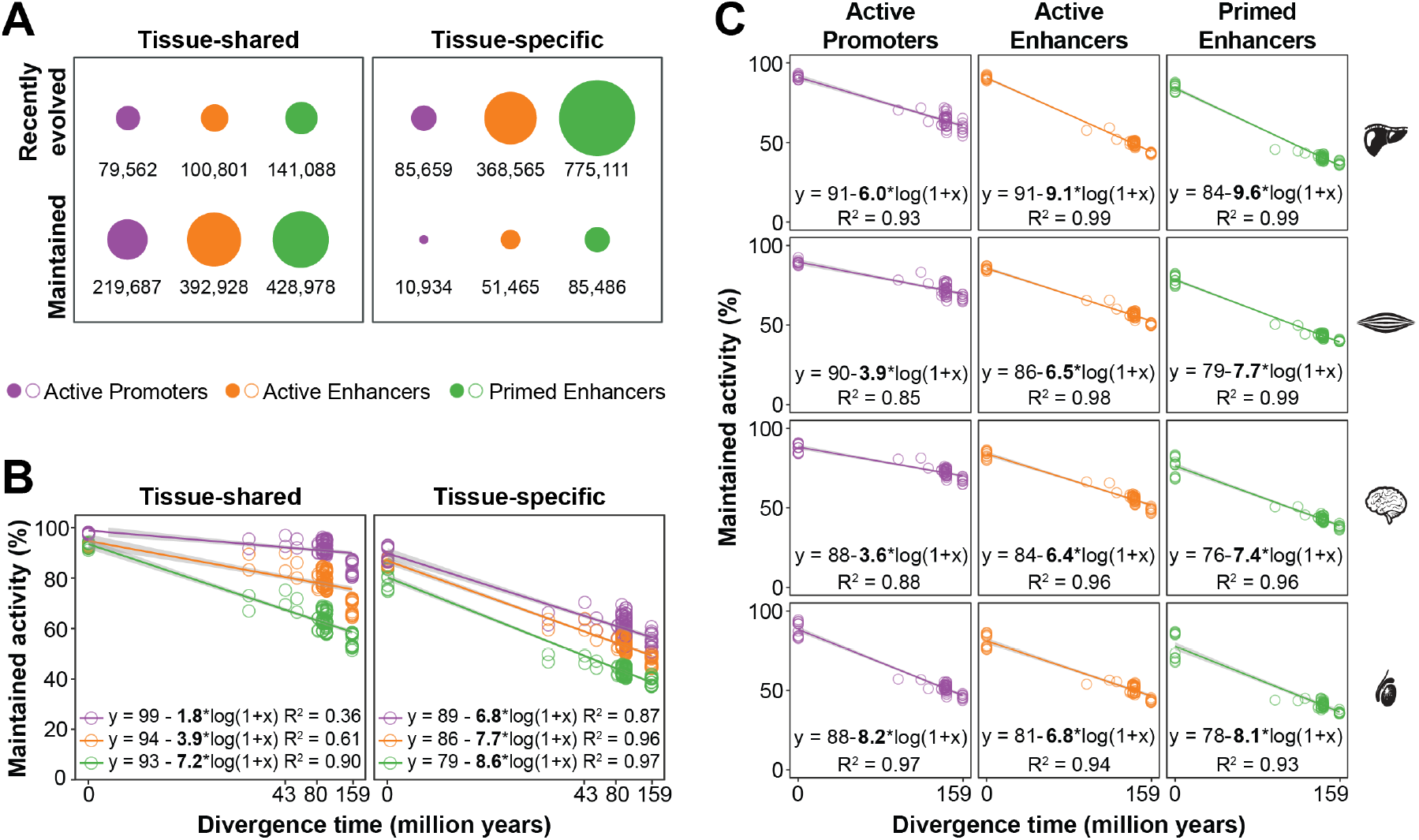
Tissue-specific regulatory regions have higher evolutionary turnover than tissue-shared regions. A) The number of tissue-shared and tissue-specific regulatory regions that are either maintained or recently evolved across all ten species (see Figure S3A). The majority of tissue-shared regulatory regions are maintained across species: 73% of active promoters, 80% of active enhancers, and 75% of primed enhancers. The majority of tissue-specific regions are recently-evolved, although 11% of active promoters, 12% of active enhancers and 10% of primed enhancers are maintained. B) Evolutionary rates of alignable tissue-shared and tissue-specific regulatory regions estimated by linear regression of activity maintenance between all pairs of species and zero points estimated from interindividual variation (Methods). For tissue-shared regions, the slope of the regression line for active promoters is lower than that of active enhancers or primed enhancers (Two-way ANOVA of linear regression: active promoters vs active enhancers p-value 0.0063; active promoters vs primed enhancers p-value 0.0056). For all tissue-specific regions, the rates of evolution are either indistinguishable or greater than that for tissue-specific primed enhancers (Two-way ANOVA of linear regression: active promoters slope vs primed enhancer slope, p-value 0.011; active enhancers slope vs primed enhancers slope, p-value 0.021). C) Evolutionary rates of tissue-specific regulatory regions further stratified by tissue of activity. The slope of the regression line for testis-specific active promoters is significantly higher than for promoters with activity specific to the liver, muscle, or brain (Two-way ANOVA of linear regression: testis-specific active promoters vs all other tissue-specific active promoters p-value 3×10^−8^). However, all tissue-specific promoters evolve more rapidly than tissue-shared promoters, regardless of their tissue of activity (Two-way ANOVA of linear regression: all tissue-specific active promoters (Figure 3B) vs tissue-shared active promoters (Figure 3B) p-value 4×10^−8^).

We quantified evolutionary rates for tissue-shared and tissue-specific regulatory regions. Through pairwise comparisons of alignable regulatory regions, we calculated the fraction of promoters and enhancers maintained between each pair of species, then used the slope of a linear fit to estimate the evolutionary rate of change (Figure 3B, Methods). Studies in single mammalian tissues have found that enhancers evolve more rapidly than promoters, but without differentiating between active and primed enhancers (Cheng et al., 2014; Villar et al., 2015). Here, we demonstrate that primed enhancers evolve much faster than active enhancers for both tissue-shared and tissue-specific regulatory elements. More importantly, we consistently found that tissue-specific regulatory regions evolved more rapidly than their tissue-shared counterparts (Figure 3B). Interestingly, tissue-specific active promoters evolved at rates comparable to enhancers, which may partly explain previous observations of fast rates of transcription start site evolution (Vierstra et al., 2014; Young et al., 2015).

We then asked whether regulatory regions evolve faster in particular tissues (Figure 3C). Among promoters, those with testis-specific activity evolved most quickly, followed by liver-specific ones. In contrast, among both active and primed enhancers, those with liver-specific activity were the fastest evolving. Brain-specific regulatory regions evolved the most slowly (Figure 3C). These tissue-specific rates of regulatory evolution give new insight into previously reported gene expression evolution rates, which found relatively small changes in brain and accelerated evolution in testis and liver (Brawand et al., 2011; Cardoso-Moreira et al., 2019). Our results suggest that both enhancers and promoters underlie the previously observed evolutionary rates of gene expression across tissues.

In sum, tissue-shared regulatory activity is a trait predictive of slower evolutionary turnover, regardless of the class of regulatory region or tissue of activity.

### Regions that switch promoter and enhancer signatures within a species are uncommon, and are not evolutionarily maintained

Prior studies have identified genomic regions that can act as either promoters or enhancers in different contexts (Dao et al., 2017), but have not evaluated the evolutionary turnover and maintenance of such dynamic regulatory regions. We defined intra-species dynamic regulatory regions as those with differing histone modification signatures across tissues in a single species (Figures S3A, 4A, Methods). Specifically, regions identified as active promoters in one tissue and active and/or primed enhancers in another tissue were defined as intra-species dynamic promoter/enhancers (dynamic P/Es). Similarly, regions identified as active enhancers in one tissue and primed enhancers in another were defined as intra-species dynamic enhancers (dynamic Es). Between the four tissues, only a small portion of each species’ regulome was intra-species dynamic on average: 7% of active promoters, 11% of active enhancers, and 7% of primed enhancers (Figure 4A).

**Figure 4:**
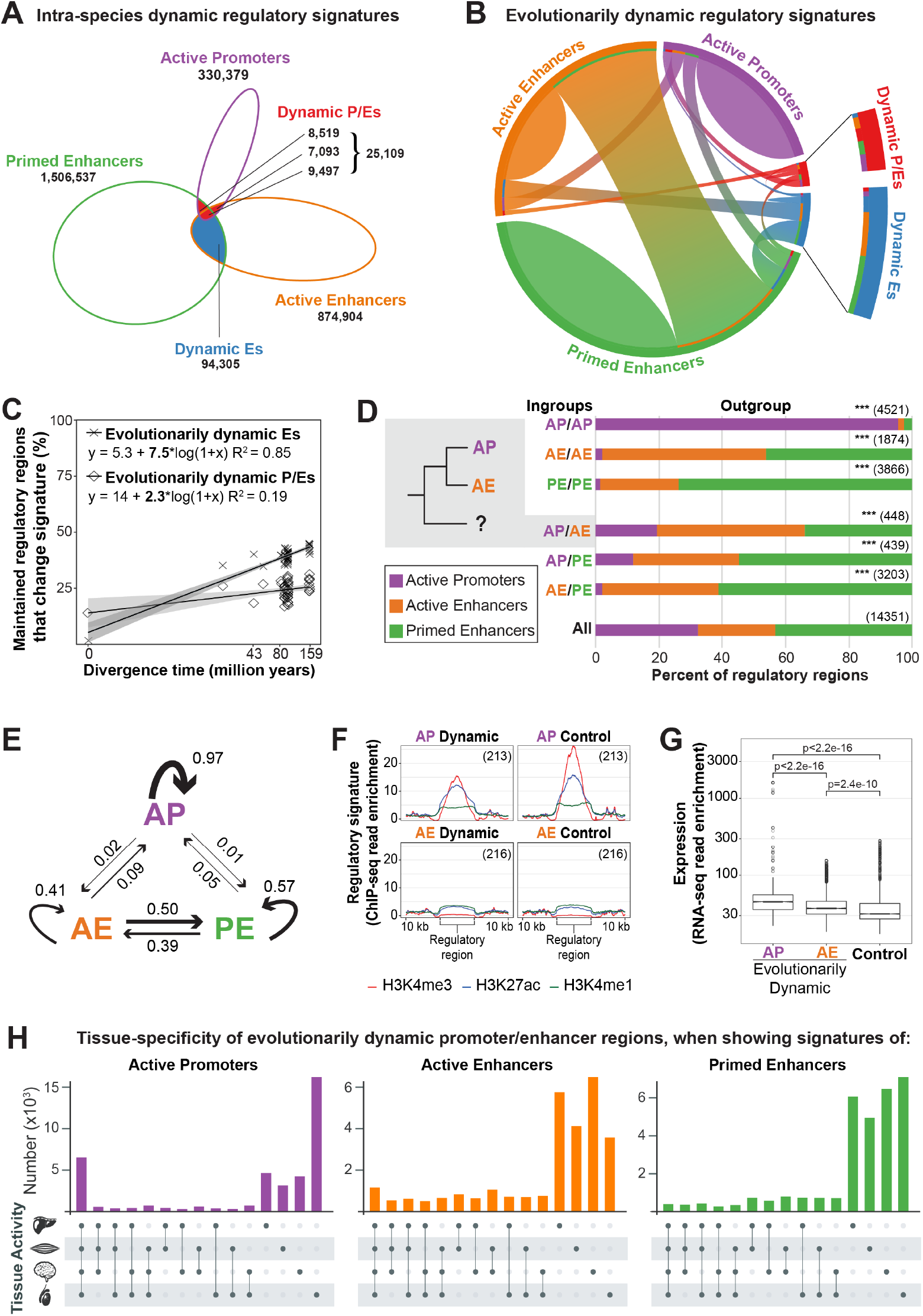
Promoter and enhancer signature is highly dynamic across species, but not within species. A) Within a species, dynamic P/Es (red) were regions identified as an active promoter in one tissue and an enhancer in another tissue, and dynamic Es (blue) were an active enhancer in one tissue and primed in another. Numbers for each category are summed across all ten species. Within a species and across tissues, only 4% of the regulome is composed of intra-species dynamic regions. B) Pairwise comparisons between maintained regulatory regions show how often regulatory signature changes between species. A substantial proportion of regulatory regions align to a region with a different regulatory signature in another species: 20% of pairwise comparisons with active promoters, 58% with active enhancers and 40% with primed enhancers are evolutionarily dynamic. Almost half of active enhancers (44%) aligned to primed enhancers. Dynamic P/Es (red) and dynamic Es (blue) almost always align to non-dynamic categories in other species (73% and 88% respectively), illustrating the evolutionary instability of this regulatory assignment. An enlargement of the intra-species dynamic regions is shown on the right for clarity. C) Evolutionary rates of changing regulatory signatures among maintained regulatory regions estimated by linear regression of pairwise comparisons. Across evolution, maintained active promoters (crosses) and active enhancers (diamonds) were more likely to change regulatory signature as evolutionary distance between species increased. D) Evolutionary directionality of dynamic regulatory signatures estimated by outgroup analysis of mouse/rat/rabbit and cat/dog/horse triads. Grey inset example: in 448 cases when a genomic region is an active promoter in one ingroup species and an active enhancer in the other, the outgroup species was most likely to be an active enhancer (46%), and least likely to be an active promoter (20%). The distributions of outgroup active promoters, active enhancers and primed enhancers for each ingroup combination is statistically different from the background (All) distribution (Chi-Square two-tailed test, *** p< 0.001). Outgroup analysis was performed separately for each triad group, and then combined (see Figure S4B and S4C). E) Composite model based on the observed likelihood of regulatory regions changing or maintaining regulatory signatures over evolution. The thickness of the lines reflects the relative likelihood of evolutionary change, as calculated from the most parsimonious evolutionary relationships from the triad data in D) and normalising the outgoing lines from each state to one. F) Validation of regulatory signature assignment using the average ChIP-seq read enrichment for evolutionarily dynamic regulatory regions and equal numbers of randomly selected control regions. Dynamic regions were the AP/AE ingroup regions identified as AE in the outgroup analysis in D. Total number of regions used are shown as insets. G) Distribution of RNA-seq read counts for evolutionarily dynamic active promoters (AP) and active enhancers (AE) shown in F, and equal numbers of randomly selected control active enhancers that are not evolutionarily dynamic (p-values calculated using one sided the t-test for greater expression). H) Tissue distribution of evolutionarily dynamic P/Es in the species where they were an active promoter, active enhancer or primed enhancer. When showing signatures of active promoters (left; purple), they were less likely to be tissue-shared and more likely to be testis-specific than all promoters (compared to Figure 2B). When showing enhancer signatures, they were more likely to be tissue-shared than all enhancers (Figure 2B bottom orange and green).

We compared the evolutionary rates of intra-species dynamic P/Es and dynamic Es with that of typical promoters and enhancers (Figure S4A, Methods). Dynamic P/Es had a higher evolutionary rate than tissue-shared active promoters or active enhancers and were more maintained than tissue-specific active promoters or active enhancers. Similarly, dynamic Es had a higher evolutionary rate than tissue-shared active enhancers and were more maintained than tissue-specific active enhancers or primed enhancers. Thus, the evolutionary stability of dynamic regulatory regions is between that of their tissue-shared and tissue-specific counterparts.

We investigated the evolutionary stability of intra-species dynamic P/Es and dynamic Es by asking how often they aligned to a dynamic region in another species (Figure 4B, Methods). The majority of intra-species dynamic P/E alignments were to non-dynamic regions in other species (73%) with approximately equal numbers aligning to active promoters, active enhancers, and primed enhancers in another species. Similarly, the majority of alignments that included intra-species dynamic Es (80%) were either to active or primed enhancers, with only 12% aligning to another dynamic E.

In sum, intra-species dynamic regions that switch promoter and enhancer signatures between tissues are relatively rare and are not maintained as intra-species dynamic regions across species.

### Promoter and enhancer signature switching is common between species

Evolution may also result in changes to the functional signatures of regulatory regions across species. Indeed, prior studies have identified a limited set of genomic regions that switch between promoter and enhancer signatures within primates or rodents (Carelli et al., 2018).

We thus investigated the evolutionary stability of histone modification signatures for all pairwise comparisons between species where regulatory activity is maintained (Figure S3A, Methods). For example, we asked how often an active promoter in mouse aligns to an active enhancer in dog – regardless of the tissue of activity. Active promoters were the most stable regulatory class across evolution: for those that were maintained as a regulatory region across species, 80% of pairwise comparisons were identified as promoters in both species (Figure 4B). Both classes of enhancers were less stable across evolution: only 42% and 60% of pairwise comparisons involving active and primed enhancers, respectively, retained the same enhancer signature between the two species.

Similar to the intra-species dynamic regions, we defined evolutionarily dynamic regions as those with different regulatory signatures between species (Figure S3A). We found that evolutionarily dynamic regions were more common than intra-species dynamic regions. For example, 15% of pairwise comparisons involving promoters were evolutionarily dynamic (Figure 4B) compared to only 7% intra-species dynamic (Figure 4A). For enhancer comparisons, 44% of active enhancers aligned to a primed enhancer in another species (Figure 4B), compared to 10% of active enhancers in one species identified as primed enhancers in a different tissue (Figure 4A). Indeed, almost half of active enhancers were readily interchangeable with primed enhancers across ten mammals, suggesting that enhancer states are in an approximate evolutionary balance.

We investigated whether regulatory regions were more likely to change signature with increasing evolutionary distance. We calculated the proportion of maintained promoters that switch between active promoter and any enhancer (evolutionarily dynamic P/Es; Figure S3A); as well as the proportion of maintained active enhancers that switch between active and primed enhancers (evolutionarily dynamic Es; Figure S3A). The proportion of maintained active promoters and active enhancers changing regulatory signature and becoming evolutionarily dynamic increased with greater evolutionary distance (Figure 4C). The rate of switching between active and primed enhancers was higher than between promoters and enhancers (Figure 4C). With this, we have quantified two evolutionary trajectories of regulatory regions: the rate of overall loss of regulatory regions (Figure 3B) and the frequency at which maintained regions change their regulatory signature across species (Figure 4C).

To examine directionality of changing regulatory signatures, we focused on species in our phylogeny with shorter evolutionary distances and clear ingroup and outgroup relationships. We separately investigated mouse and rat with rabbit as outgroup, and cat and dog with horse as outgroup. We considered only regulatory regions that were maintained across all three species. For each regulatory region, we determined the regulatory signature in the outgroup species given the signatures in the two ingroup species, regardless of the tissue of activity (Figure S3A). As expected, when a genomic region was defined as an active promoter in both ingroup species, it was also defined as an active promoter in the outgroup 95% of the time (Figure 4D, S4B, S4C).

When a genomic region was consistently identified as an active enhancer in both ingroup species, it was a primed enhancer 46% of the time in the outgroup (Figures 4D, S4B, S4C). Correspondingly, when a region was identified as a primed enhancer in both ingroup species, it was an outgroup active enhancer 25% of the time. These results further demonstrate that active and primed enhancers are readily interchangeable throughout evolution. Similarly, when a region was defined as an active enhancer in one ingroup species and a primed enhancer in the other, it was identified as an outgroup active enhancer 37% of the time and as a primed enhancer 61% of the time. This suggests that for evolutionarily dynamic Es, the ancestral state is almost twice as likely to be a primed enhancer than an active enhancer. Thus, primed enhancers are more likely to evolve into active enhancers than the reverse, yet both types of changes are widespread.

Our data enabled us to quantitatively investigate the suggested model that promoters arise from ancestral enhancers (Carelli et al., 2018). Regions identified as an active promoter in one ingroup species and an active or primed enhancer in the other were identified as an enhancer in the outgroup species more than 80% of the time (Figures 4D, S4B, S4C). The similar contribution of active and primed enhancers in the outgroup is likely due to their rapid evolutionary interchange. Overall, promoters arise from enhancers six times more often than enhancers arise from promoters.

We used the frequency of regulatory signature change observed in the outgroup analysis to model regulatory signature evolution (Figure 4E, Methods). Our model predicts that active promoters are most likely to maintain their signature, and primed enhancers are about as likely to evolve to active enhancer signatures as they are to remain primed. Active enhancers have two equally likely evolutionary fates: maintaining their signature or evolving into primed enhancers.

Finally, we validated the evolutionary switching of regulatory signatures from enhancers to promoters by examining the enrichment of histone modifications and transcription around evolutionarily dynamic P/Es identified in the outgroup analysis (Figures 4E, S4B, and S4C). We used parsimony to select regions that were most likely to represent evolutionary switches from active enhancer to active promoter (ingroups: active promoter, active enhancer; outgroup: active enhancer), and compared the ChIP-seq and RNA-seq read enrichment between regions marked as active promoters and active enhancers in the ingroups. The regions showed characteristic chromatin signatures of active enhancers and active promoters (Figure 4F). Furthermore, ingroup active promoters had increased transcription of flanking regions compared to ingroup active enhancers (Figure 4G), indicating that the regulatory signature change leads to higher transcriptional activity.

We initially defined evolutionarily dynamic P/E regions without considering their tissue of activity (Figure S3A). We examined the tissue-specificity of these regions and compared it to the overall tissue-specificity pattern for promoters and enhancers (Figure 2B). For each region, we separately characterized the tissue-specificity in species where it showed signatures of an active promoter, active enhancer, or primed enhancer (Figure 4H). In species where evolutionarily dynamic regions had an enhancer signature, they were mostly tissue-specific, similar to the trend for all enhancers (Figure 2B) and were only slightly more likely to be tissue-shared than all other enhancers (24% evolutionarily dynamic enhancers active across more than two tissues, compared to 20% of all enhancers; binomial test p-value <2.2×10^−16^). When evolutionarily dynamic regions showed promoter signatures changes to tissue-specificity were more pronounced, with only 16% of them being active across all four tissues (Figure 4H) compared to 37% of all promoters (Figure 2B; binomial test p-value < 2.2×10^−16^). Interestingly, in the species where evolutionarily dynamic P/Es had promoter signature, 41% were testis-specific (Figure 4H), which is significantly higher than the 25% observed for all promoters (Figure 2B; binomial test p-value < 2.2×10^−16^). These results suggest that evolutionarily dynamic promoters change both regulatory signature and tissue-specificity between species.

### LINEs are a versatile source of regulatory activity

We next leveraged our dataset to ask how specific classes of repeat elements contribute to the evolution of tissue-specific and tissue-shared regulatory activity across mammals. We separately analysed recently-evolved and maintained regulatory regions (Figure 3A) and identified which transposable elements they overlap. We grouped transposable elements into LINEs, SINEs, LTRs and DNA transposons. Within each of these groups, we compared the enrichment of annotated transposable element families between tissue-specific and tissue-shared regulatory regions (Methods). Various families of transposable elements within the LTR and SINE groups such as Alu, B2 and ERVL elements contributed to tissue-specific and tissue-shared active promoters in a lineage-specific manner (Figures 5A and S5A), in line with previous observations (Franke et al., 2017; Trizzino et al., 2018).

Strikingly, across all study species we found that tissue-specific active promoters were enriched with LINE L1 family of transposons as compared to their tissue-shared counterparts, and tissue-shared active promoters were enriched in the LINE L2 family (Figures 5A and S5A). This was observed for both recently-evolved (Figure 5A) and maintained (Figure S5A) active promoters. The same trend of LINE L1 and L2 enrichment was observed in recently-evolved and maintained active enhancers, although the trend is weaker, and was not as evident for primed enhancers (Figures 5A and S5A).

**Figure 5:**
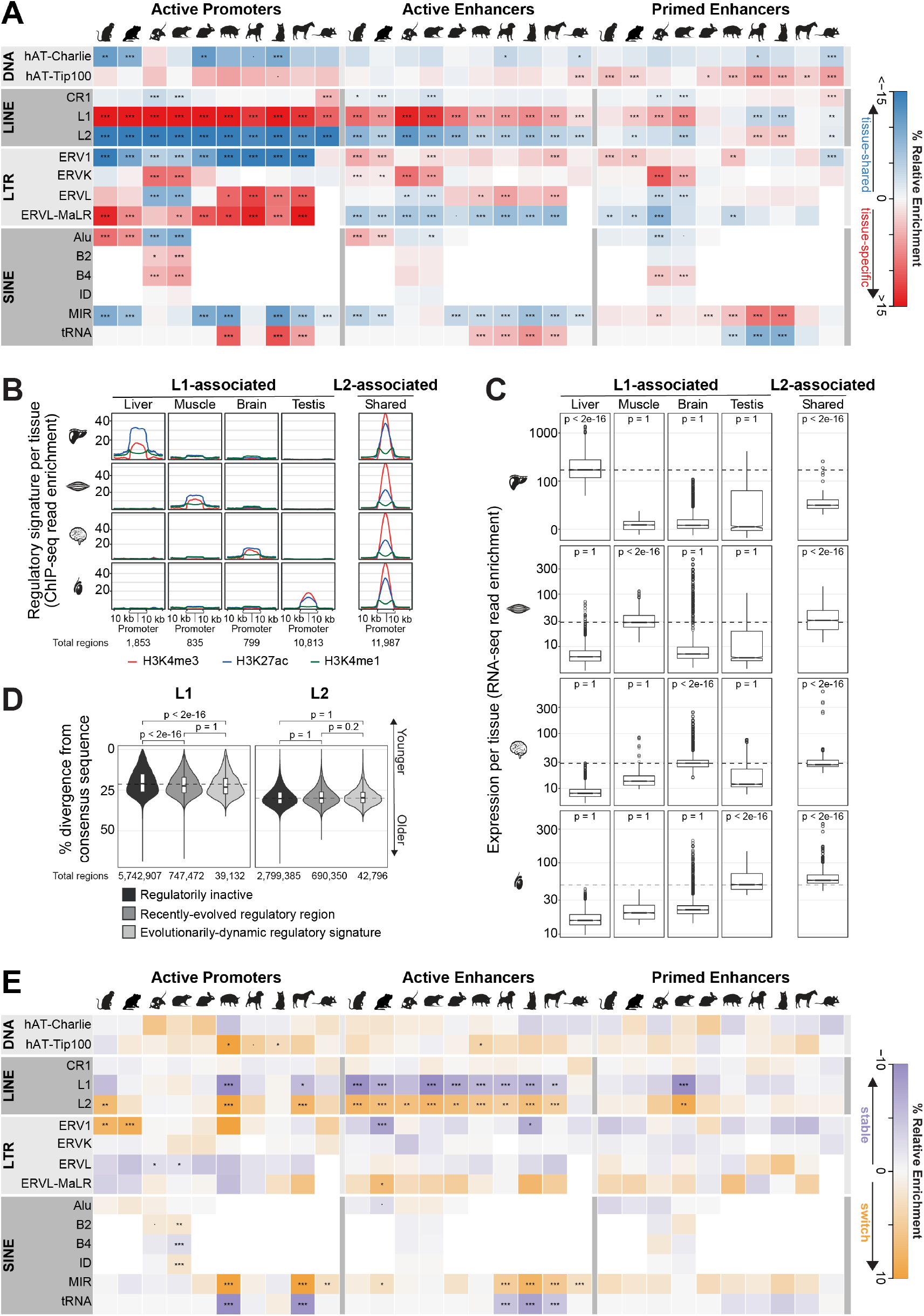
Distinct families of repetitive elements contribute to recently-evolved and maintained regulatory regions. A) Relative enrichment of recently-evolved tissue-shared versus tissue-specific regulatory regions for selected transposable element families shown as a heatmap. Within each family, significance of tissue-specific vs. tissue-share proportions calculated with the Z-test and Bonferroni correction (P-values *** < 0.001; ** < 0.01; * < 0.05; - < 0.1. See Figure S5A for maintained regions). B) Validation of tissue-specific activity using the average ChIP-seq read enrichment for recently-evolved active promoters associated with LINE L1s and L2s and their flanking regions. C) Distribution of RNA-seq read counts for the promoter flanking regions in B). Dotted lines represent the median of tissue-specific RNA-seq enrichment for the tissue profiled. P-values calculated using one sided the Wilcoxon test; within row test if read counts in each LINE-associated region type (column) is greater than in all other regions. D) Estimated age of LINE L1 and L2 elements, as inferred by the number of substitutions from consensus sequence. LINE L1s that overlap regulatory regions (medium and light grey) are significantly older than inactive L1s (dark grey), while regulatorily active L2 elements are of similar age to inactive L2s. Dotted line is the median % divergence of the corresponding regulatorily inactive LINEs and p-values calculated using one sided the Wilcoxon test for greater sequence divergence. Divergence is shown for all ten species combined; see Figure S5B for per-species divergences. E) Heatmap of relative enrichment in transposable element families for regulatory regions with evolutionarily dynamic (switch) versus stable signatures. Within each family, significance of evolutionarily dynamic vs. stable proportions calculated with the Z-test and Bonferroni correction (P-values *** < 0.001; ** < 0.01; * < 0.05; - < 0.1).

To gain insight into the transcriptional consequences of LINEs, we examined the histone modification enrichments (Figure 5B) and gene expression (Figure 5C) within 10 Kb of active regulatory regions overlapping LINE elements. Among recently evolved active promoters that were both tissue-shared and contained L2 elements (7% of recently evolved promoters), all had high enrichment of H3K4me3 and H3K27ac and increased nearby transcription. The 9% of the recently evolved active promoters that were both tissue-specific and contained L1 elements showed enrichment only in the relevant tissue.

We compared the transposable element enrichment in evolutionarily dynamic P/E regions to those regions that retain stable regulatory signatures between species (Figure 5E, Methods). Among active enhancers, evolutionarily dynamic P/Es showed relative enrichment in the LINE L2 family compared to stable active enhancers. In contrast, stable active enhancers were enriched for the LINE L1 family. This trend is also evident for active promoters and primed enhancers in some lineages.

We investigated whether regulatory activity was associated with the evolutionary timing of LINE retrotransposition. The age of each LINE was estimated by its divergence from the consensus sequence. LINEs were divided into those that overlap identified regulatory regions and those that do not (Figure 5D, Figure S5B). As expected, LINE L2 elements were older than L1 elements regardless of regulatory association (Chalopin et al., 2015). For all study species, the age of LINE L2s was similar for recently evolved tissue-shared regulatory regions, evolutionarily dynamic regions, and for L2s not associated with any regulatory activity.

LINE L1s that overlapped regulatory regions were significantly more diverged (Figure S5D) and thus older than those that were not regulatorily active. Specifically, regulatory regions that overlapped L1s were less likely to overlap the youngest L1 elements. This effect varied across species, and was especially pronounced in rodents, primates, and opossum, where many L1 elements arose recently and have remained regulatorily inactive (Figure S5B). Using the reported mutation rates of primate LINEs (Thybert et al., 2018), we estimated that the expansion of L2s happened before the split of eutherian mammals (~100 million years ago), and the L1 expansion after the split, consistent with previous whole genome findings (Lovsin et al., 2001). To find evidence of selection in LINEs, we compared sequence constraint between tissue-specific regulatory regions overlapping L1s and L2s, and found that those overlapping L2s are significantly more constrained across their whole length (Figure S5C) and contain a larger number of constrained elements (Figure S5D). This suggests that lineage-specific genetic variation unmasks latent regulatory potential in existing LINE L2s.

Across the mammalian lineage, active regulatory regions consistently associated with LINE L1 transposable elements if they were tissue-specific, and with LINE L2s if they were tissue-shared (Figures 5A and S5A). LINE L2s also consistently associated with evolutionarily dynamic regulatory regions (Figure 5E), which frequently change both regulatory signature and tissue of activity, suggesting that LINE L2s provide a more versatile potential for transcriptional regulation than LINE L1s.

These analyses demonstrate the contribution of LINEs in shaping gene regulatory landscapes across the mammalian regulome.

## DISCUSSION

Regulatory landscapes are composed of tissue-specific and tissue-shared regions that appear complex and evolutionarily unstable. We have created a comprehensive experimental dataset characterising how tissue-specific transcriptional regulation has evolved from a common mammalian ancestor 159 million years ago. Using four adult primary tissues from ten species, we identified nearly 3 million regulatory regions, and quantified the associated gene expression. Our analyses have given high-resolution insight into the evolutionary relationship between tissue-specificity and functional maintenance, characterized changing regulatory signatures across tissues and species, and revealed how LINE transposable elements evolutionarily shape tissue-specificity.

Our analyses of the mechanisms of regulatory evolution between species and tissues have limitations. First, the four tissues we profiled do not represent all possible cell types, though the distinctive evolutionary mechanisms we have identified are likely robust, because our categorization of tissue-shared or tissue-specific is unlikely to substantially change with the addition of more cell types (Cheng et al., 2014). Second, our analysis does not capture all enhancers and promoters; like every method to define regulatory regions, it has specific advantages, disadvantages, and biases (Andersson and Sandelin, 2020). We used a widely-employed approach of combining three histone modification and performed all experiments in at least biological triplicates, yet this strategy cannot identify alternative promoters at high resolution as can techniques like CAGE (Forrest et al., 2014). Furthermore, H3K4me1, which differentiates active and primed enhancers, is more variable between replicates than other histone marks (Figure S2A). Third, to fully explore how tissue-specific and tissue-shared regulomes interact to shape the evolution of gene expression would require quantifying three-dimensional contacts.

### Regulatory roles change readily across evolution

Our results reveal that primed and active enhancers are frequently redeployed across evolution into different regulatory roles. Within a species, only a small subset of promoters interchange regulatory roles with enhancers, in line with previous studies (Dao et al., 2017; Leung et al., 2015). Between species, there was suggestive evidence that ancestral enhancers can evolve to promoters in somatic tissues (Carelli et al., 2018). By analysing a large number of species, characterizing a greater diversity of regulatory regions, and including a germline tissue, we discovered that changing regulatory roles is, in fact, a frequent event in mammalian evolution. One-fifth of alignments with maintained promoters and almost half of alignments with enhancers showed evidence of such interchange between species. The observed frequent evolutionary interchange of active and primed enhancers may be the result of a birth-death balance, or potentially reflect a plasticity in the histone signatures of enhancers. We demonstrated that enhancers interchange regulatory signatures with promoters across evolution, most frequently in the testis. The distinct regulatory plasticity in testis supports a model wherein germline tissues have unique roles in evolution.

### LINE transposable elements shape regulatory evolution across mammals

One of our most striking results is the opposing contributions of LINE L1s and L2s to regulatory evolution. Regulatorily active LINEs do not generally arise from lineage-specific insertions, suggesting that the predominant mechanisms for regulatory activation is mutation of ancient elements – even for those with lineage specific activity. Multiple studies have characterized the contribution of lineage-specific insertions of transposable elements to regulatory evolution (Cao et al., 2019; Chuong et al., 2016; Jacques et al., 2013; Trizzino et al., 2017). In contrast, the regulatory potential of more ancient insertions of transposable elements has been less studied (Simonti et al., 2017; Trizzino et al., 2018). LINE L1s are transcribed in a cell-type specific manner (Philippe et al., 2016), which corresponds to our findings that L1s are associated with tissue-specific regulatory activity. LINE L2s have been less studied, though recently shown to be ubiquitously expressed as miRNAs (Petri et al., 2019) and to have promoter and enhancer activity in human tissues (Cao et al., 2019). Our data showed that LINEs, both L2s and L1s, are widely used across mammals as an evolutionary substrate for new promoter and enhancer regulatory activity. LINE L2s are associated with tissue-shared regulatory activity and evolutionarily dynamic promoter/enhancers. LINE L1s, in contrast, are associated with tissue-specific regulatory regions, as well as those with stable regulatory signatures that do not switch between promoter and enhancer regulatory signatures.

By mapping the dynamic mammalian regulome across ten species, we reveal the complex, evolutionarily unstable regulatory landscapes underpinning stable tissue phenotypes and a role for ancient mammalian repeats in shaping their plasticity.

## ETHICS STATEMENT

The investigation was approved by the Animal Welfare and Ethics Review Board, under reference number NRWF-DO-02v2, following the Cancer Research UK Cambridge Institute guidelines on use of animals in experimental studies.

## AUTHOR CONTRIBUTIONS

M.R., E.S., D.T.O. and P.F. designed experiments; E.S., D.V. and A.R. performed experiments: M.R., O.I., F.M., E.S. and R.R. analysed the data; M.R., E.S., D.T.O. and P.F. wrote the manuscript; D.T.O. and P.F. oversaw the work. All authors read and approved the final manuscript.

## ACKNOWLEDGEMENTS

We thank Camille Berthelot for important insight that helped define the focus of the work and feedback on the manuscript; Vasavi Sundaram for help with analysis software and careful reading of the manuscript; Sarah Elderkin, Elissavet Kentepozidou, Simen R. Sandve, Martina Rimoldi and Emily Wong for critical reading of the manuscript; Francesco Nicola Carelli and Anne-Ruxandra Carvunis for helpful discussions; Margus Lukk, Tim Rayner, Gordon Brown, Richard Bowers for data management; Matthew Clayton and Mike Mitchell and others from the CRUK-CI Biological Resources Unit, Genomics, and Bioinformatics cores for technical assistance; James Turner and Bryony Leeke from the Francis Crick Institute; Fernando Montesso at the Animal Health Trust in Newmarket; David Farningham and staff at the MRC Harwell Centre for Macaques; Colin Windle and staff at the University Biomedical Service in Cambridge; and Toni Nieto, Gloria Nejar, and Toni Bermúdez from Isoquimen, S.L. in Barcelona for help with acquiring tissues. We thank the staff at Dstl. We acknowledge Spencer Phillips for producing species and tissue images. Finally, we would like to give acknowledgement to the animals used in this study.

Funding for this study was provided by Wellcome (WT108749/Z/15/Z, WT202878/Z/16/Z, WT202878/B/16/Z), European Research Council (615584, 788937), Cancer Research UK (20412), Helmholtz Society, and the European Molecular Biology Laboratory.

## SUPPLEMENTAL FIGURES AND LEGENDS

**Figure S1:**
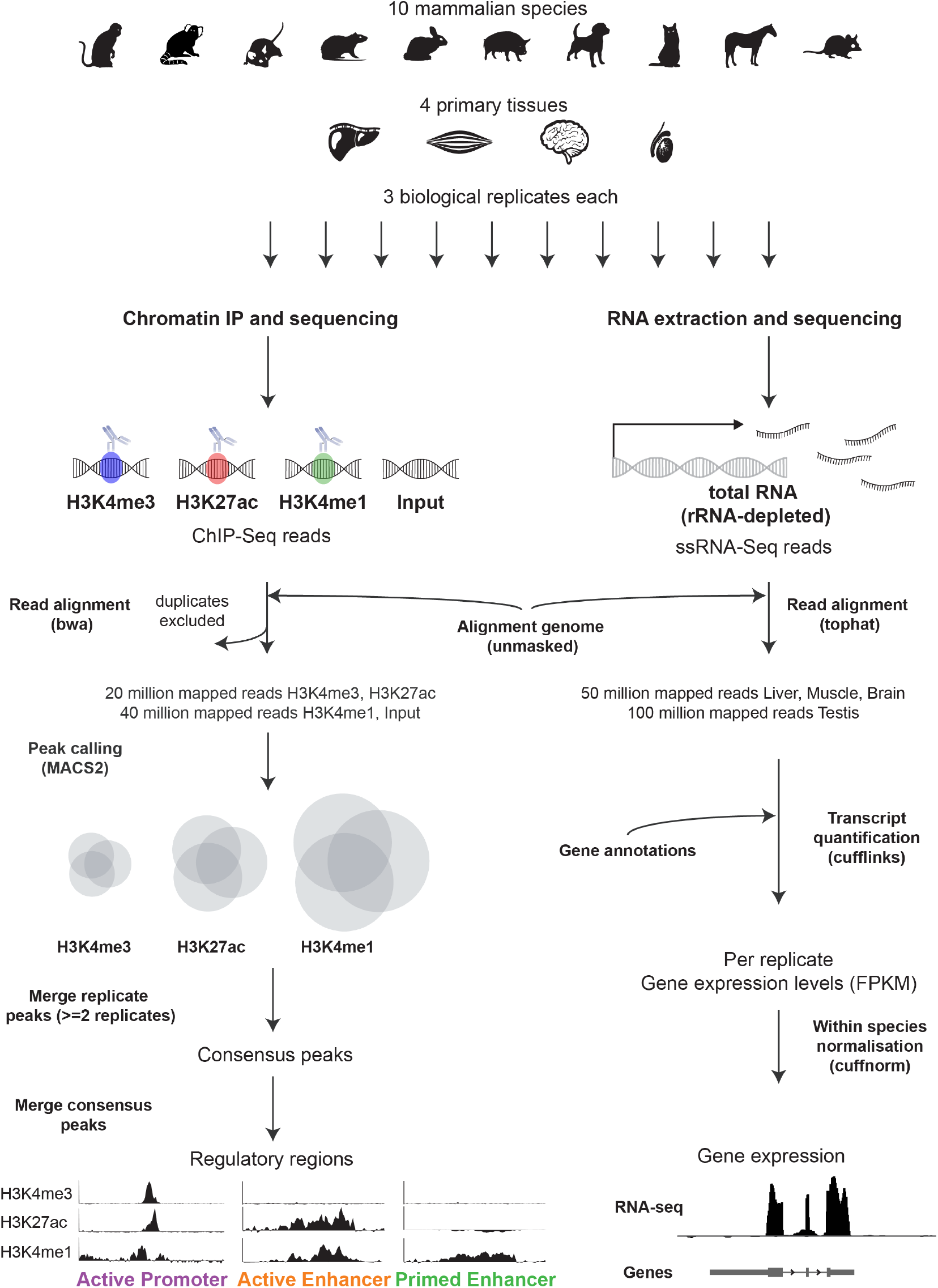
Experimental Overview. Functional genomics experiments were performed on 10 mammalian species (macaque, marmoset, mouse, rat, rabbit, pig, dog, cat, horse, opossum) and 4 primary adult tissues (liver, muscle, brain, and testis), with 3 biological replicates (individuals) for each. ChIP-seq for 3 different histone modifications (H3K4me3, H3K27ac, and H3K4me1) was used these to identify regulatory regions genome wide (active promoters, active enhancers, and primed enhancers). Within each biological replicate, ChIP-seq and input libraries were mapped, duplicate reads were removed, but multi-maping reads were retained to aid mapping across transposable elements, and then randomly subsampled to the same depth before peak calling. For each histone modification, only those peaks present in at least two replicates (consensus) were merged and kept for further analyses. Finally, regulatory regions were defined from the overlap of consensus peaks for the three histone modifications. Total RNA-seq was performed on matched tissue samples and mapped to the respective genome. Within each biological replicate, RNA-seq libraries were subsampled to the same depth per tissue. The subsampled sets were used to quantify gene expression levels within each replicate, and these were normalized within each species. (See Figure 1 for data overview.)

**Figure S2:**
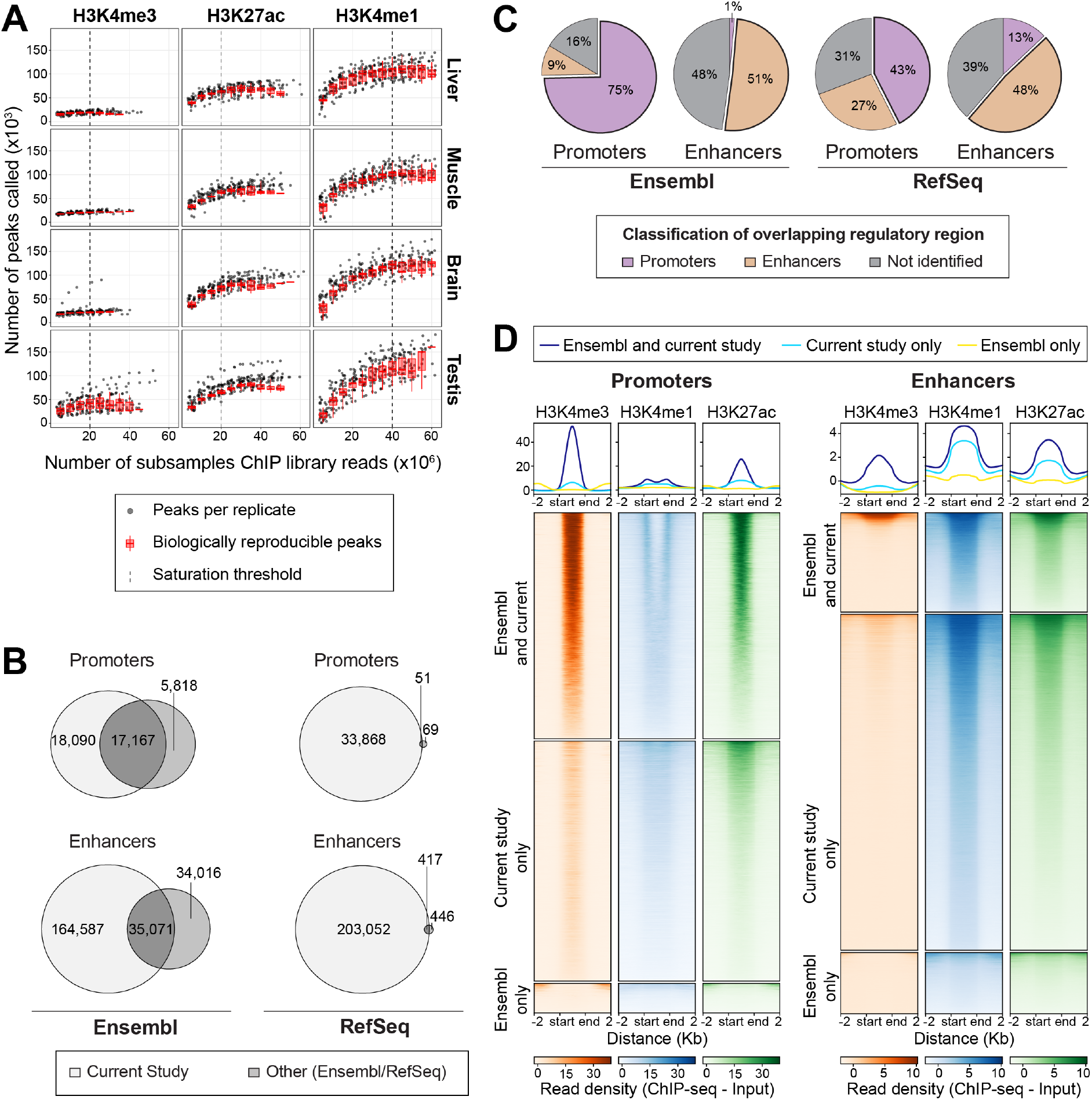
Validation of ChIP-seq and regulatory regions in Mouse (See also Figure 1). A) To choose an appropriate subsampling threshold for all species, tissues and ChIP-seq libraries were randomly subsampled across all replicates, and peaks per replicate called. Within each species and tissue, we also calculated the number of consensus regions called for each histone modification. H3K4me3 and H3K27ac signal saturated at 20 million reads in all tissues, while 40 million reads were needed to saturate H3K4me1 signal. ChIP-seq signal reached saturation in all species and tissues. B) Venn diagrams showing overlap between: (left) the promoters and enhancers called in the current study and promoters and enhancers in the Ensembl regulatory build (Zerbino et al., 2016),); and (right) promoters and enhancers experimentally validated and listed in RefSeq (O’Leary et al., 2016). For this analysis, we grouped active and primed enhancers together to make them comparable to the previous studies. The regulatory landscape of four mouse tissues from the current study recovers 75% of known mouse promoters and enhancers. C) Further details on regulatory regions from external databases shown in B). The recovered regulatory regions are mostly consistently called as the same regulatory type in external datasets and our own data. 76% of Ensembl promoters and 84% of Ensembl enhancers are consistent with our own calls, while the agreement with RefSeq is 56%of for promoters and 63% RefSeq for enhancers. D) We compared the ChIP-seq read enrichments for promoters and enhancers that were uniquely identified here (Current study only, panel B) to those that overlap regulatory regions called in the Ensembl regulatory build (Ensembl and current study, panel B), and those called in the Ensembl regulatory build but not identified here (Ensembl only, panel B). The top panel shows density plots of average fold enrichment (over input) for all regions in each category for all regions in each category; the bottom panel shows fold enrichment for each regulatory region. The length of each regulatory region was normalized, as indicated by the start and end, and windows 2 Kb upstream and downstream of regulatory regions are also shown.

**Figure S3:**
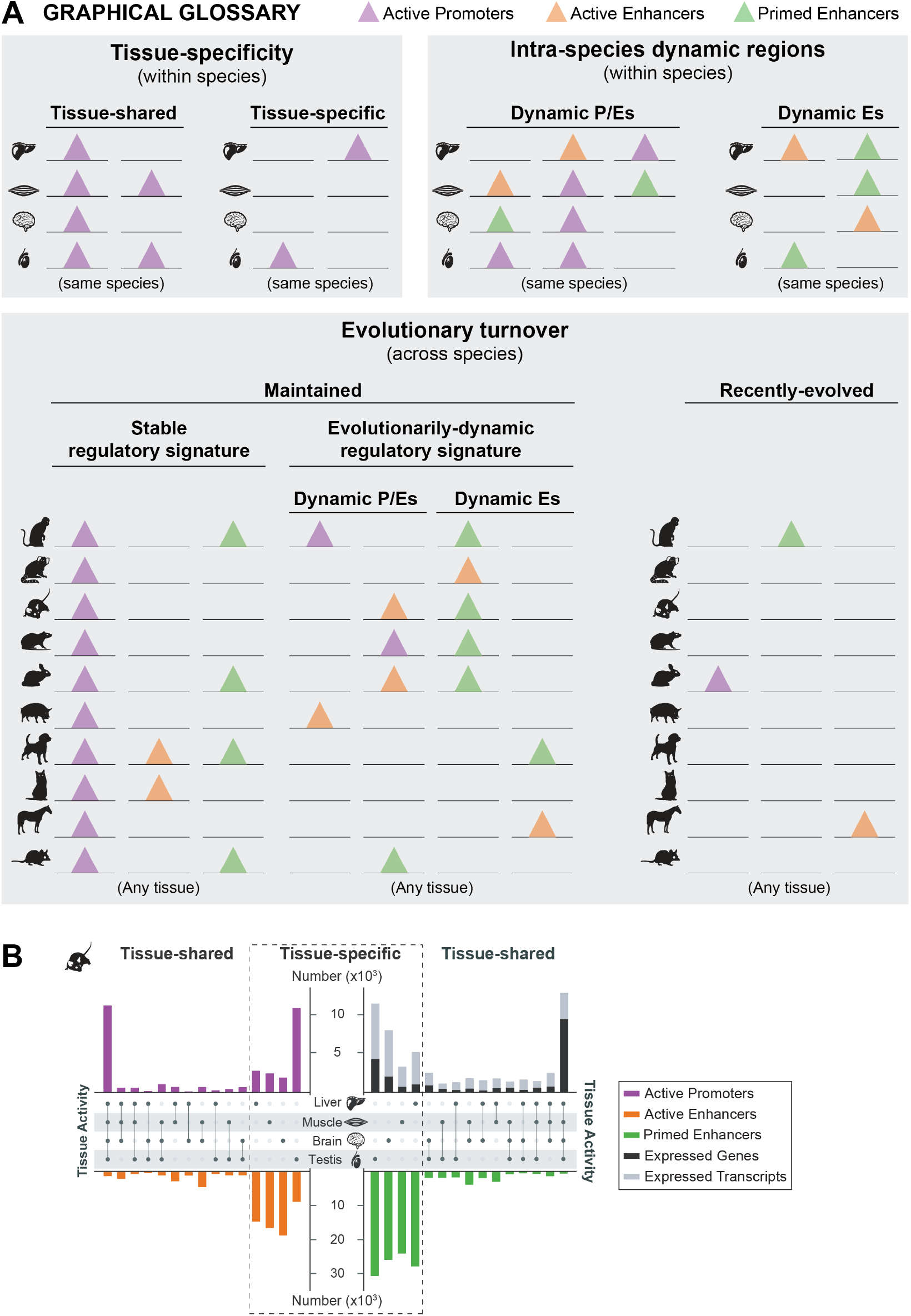
Tissue-specificity of regulatory regions and expressed genes in mouse. A) Graphical glossary showing cartoon examples of how regulatory regions were categorized within a species (top panels) and across species (bottom panel). Stacked lines indicate the same genomic region across tissues (top panels) or aligned genomic regions across species (bottom panel). Triangles indicate a regulatory assignment of active promoter (purple), active enhancer (orange), or primed enhancer (green). Specifically, within a species regulatory regions were defined as either **tissue-shared** if they were identified as a regulatory region with the same signature (i.e. active promoter, active enhancer, or primed enhancer) in two or more tissues; or **tissue-specific** if they were identified as a regulatory region in only one of the four study tissues. Within a species, regulatory regions were defined as **intra-species dynamic** regions if they were identified as one type of regulatory region in one tissue, and another type of regulatory region in another tissue of the same species. Intra-species dynamic promoter/enhancers (**dynamic P/Es**) were defined as any genomic region identified as an active promoter in at least one of the four tissues, and an enhancer (active and/or primed) in at least one other tissue of the same species. Similarly, intra-species dynamic enhancers (**dynamic Es**) were defined as any genomic region identified as an active enhancer in one or more tissues and as a primed enhancer in at least one other tissue of the same species, but not if it was also found to be an active promoter in a tissue. By aligning genomic regions across species, regulatory regions were identified as either **maintained** (identified as a regulatory region in two or more species, regardless of the type of regulatory region or tissue of activity) or **recently-evolved** (identified as a regulatory region in only one of the ten study species). Maintained regions were further classified either as those with a **stable regulatory signature** (identified as the same type of regulatory region in all species where regulatorily active) or those with an **evolutionarily dynamic regulatory signature** (identified as one type of regulatory region in one or more species, and another type of regulatory region in at least one other species). Evolutionarily dynamic regulatory regions were defined as either **evolutionarily dynamic P/Es** (identified as a promoter in at least one species and an active and/or primed enhancer in at least one other species) or **evolutionarily dynamic Es** (identified as an active enhancer in at least one species and a primed enhancer in at least one other species, but never an active promoter). Multiple possible examples are shown for each regulatory region category, however these are not exclusive, as many additional scenarios are possible. B) Tissue-specificity of regulatory regions and gene expression is shown for mouse. The ratio of tissue-specific transcripts compared to genes expressed is higher than the ratio of tissue-shared transcripts to genes expressed. Analysis is the same as for Figure 2B, which shows combined values for all ten study species.

**Figure S4:**
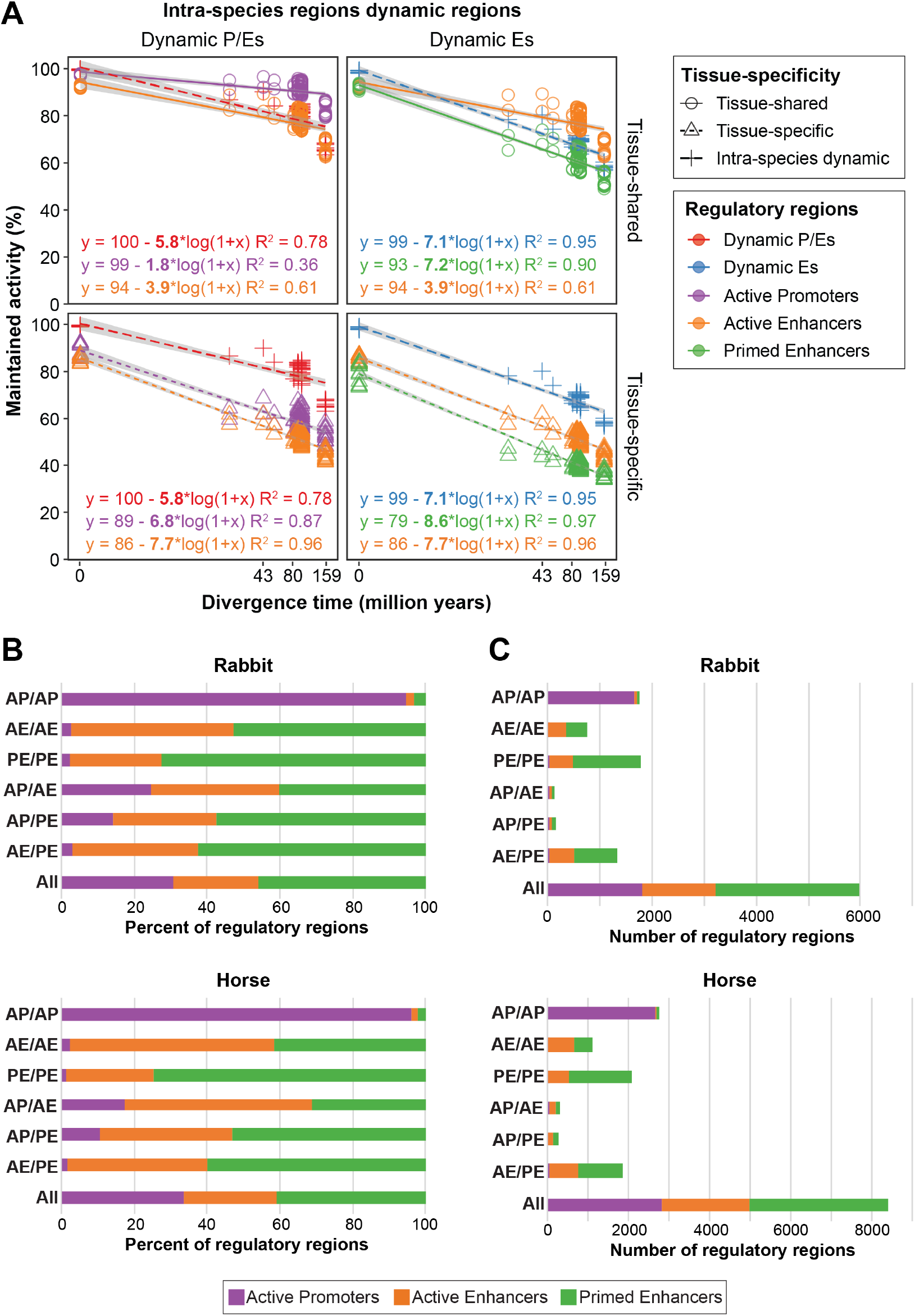
Evolutionary turnover of intra-species dynamic and evolutionarily dynamic regulatory regions. A) Similar to Figure 3B, we determined the functional conservation of intra-species dynamic regulatory regions. We compared evolutionarily dynamic promoters (red) to tissue-shared (top left) and tissue-specific (bottom left) active promoters and enhancers, and found that their evolutionary rate is intermediate between the rate of tissue-shared active promoters and enhancers. We also compared evolutionarily dynamic enhancers (blue) to tissue-shared (top right) and tissue-specific (bottom right) active and primed enhancers, and found that the rates are intermediate between tissue-shared active and primed enhancers. Evolutionary turnover rates were estimated by linear regression of activity conservation between all pairs of species for the evolutionarily dynamic regions. B) Outgroup analysis of triad species, showing results separately for mouse/rat/rabbit and cat/dog/horse (combined results are shown in Figure 4D). Given the combination of regulatory signatures in the ingroups (cat/dog or mouse/rat), we tested the signature in the outgroup species (horse or rabbit, respectively) to assay the directionality of the evolutionary change. The background distribution (All) corresponds to all regulatory regions maintained across all three species. Active enhancers often evolve from primed enhancers, and vice versa. Active promoters are more stable, but when a genomic region does change between active promoter in one species and enhancer in another, the direction is more often from enhancer to promoter. Values are shown as percentages. C) Same as in B, but with values shown as raw numbers, rather than percentages.

**Figure S5:**
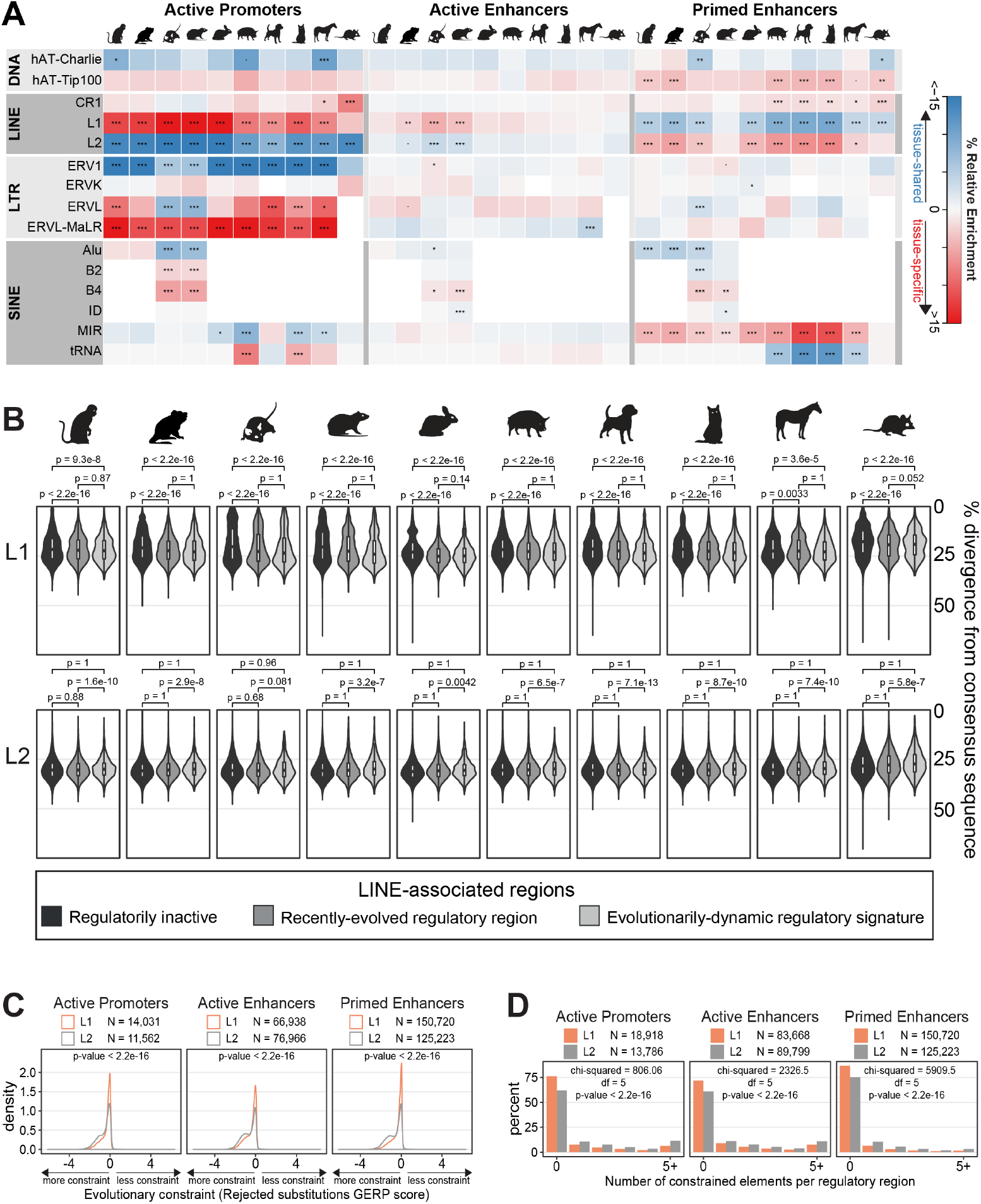
Regulatory active LINE L1s are not represented in the most recent whole genome expansions. (Related to Figure 5). A) The relative enrichment of maintained tissue-shared versus tissue-specific regulatory regions for transposable element families is shown as a heatmap [red=enriched in tissue-specific; blue=enriched in tissue-shared; white = comparable contribution to tissue-specific and tissue-shared]. LINE L1s are enriched in tissue-specific regulatory regions, while LINE L2s are enriched in tissue-shared. Within each family, significance of tissue-specific vs. tissue-share proportions calculated with the Z-test and Bonferroni correction (P-values *** < 0.001; ** < 0.01; * < 0.05; - < 0.1). A similar heatmap for recently-evolved tissue-shared versus tissue-specific regulatory regions is shown in Figure 5A). B) For every regulatory active (medium and light greys) and inactive (dark grey) LINE L1 and L2 within each species, we calculated the number of mutations from consensus sequence as a proxy for transposable element age. Analysis is the same as for Figure 5D, which showed divergence for all ten species combined. p-values were calculated using one sided the Wilcoxon test for greater sequence divergence. C) The distribution of GERP scores for alignable tissue-specific regulatory regions associated with LINE L1s and L2s. A negative GERP scores indicates more rejected substitutions than expected and is evidence of evolutionary selection. LINE L2s, even when tissue-specific in activity, are more constrained than their L1 associated tissue-specific counterparts. P-values calculated using the Wilcoxon test. D) Distribution of the percent of tissue-specific regulatory regions associated with LINE L1s and L2s according to the total counts of constrained elements they contain, regardless of alignability. Tissue-specific LINE L2s less commonly have no constrained elements, and more commonly have 1 or more constrained elements than LINE L1s. The Chi-square test was performed on raw counts of total LINE L1 and L2 regions.

## SUPPLEMENTAL TABLES AND LEGENDS

**Table S1:**
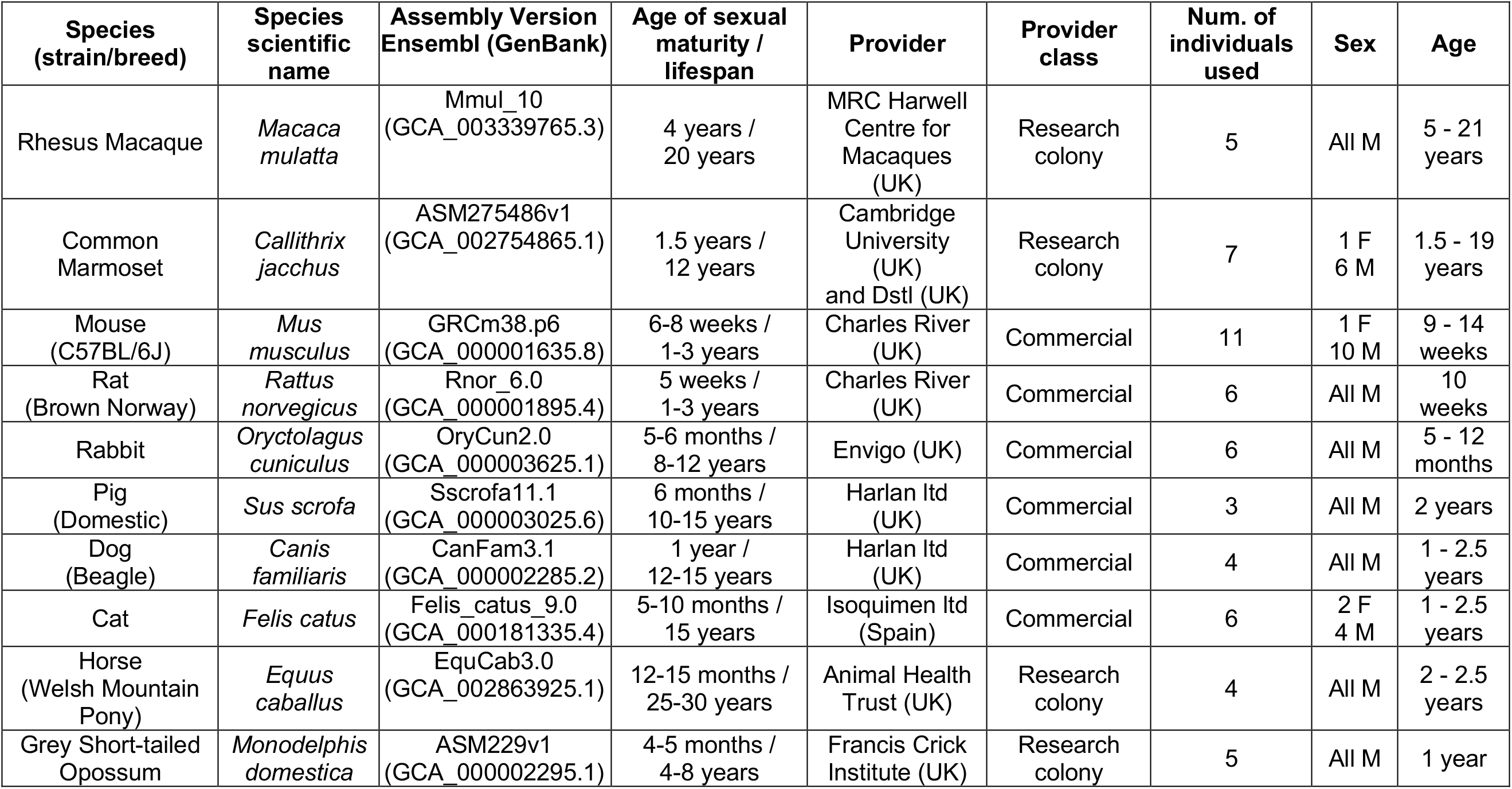
Species used in this study. Overview of the experimental samples and genomic versions used in the study, for more details see also **Table S2**.

| Species (strain/breed) | Species scientific name | Assembly Version Ensembl (GenBank) | Age of sexual maturity / lifespan | Provider | Provider class | Num. of individuals used | Sex | Age |
| --- | --- | --- | --- | --- | --- | --- | --- | --- |
| Rhesus Macaque | *Macaca mulatta* | Mmul_10 (GCA_003339765.3) | 4 years / 20 years | MRC Harwell Centre for Macaques (UK) | Research colony | 5 | All M | 5 - 21 years |
| Common Marmoset | *Callithrix jacchus* | ASM275486v1 (GCA_002754865.1) | 1.5 years / 12 years | Cambridge University (UK) and Dstl (UK) | Research colony | 7 | 1 F<br>6 M | 1.5 - 19 years |
| Mouse (C57BL/6J) | *Mus musculus* | GRCm38.p6 (GCA_000001635.8) | 6-8 weeks / 1-3 years | Charles River (UK) | Commercial | 11 | 1 F<br>10 M | 9 - 14 weeks |
| Rat (Brown Norway) | *Rattus norvegicus* | Rnor_6.0 (GCA_000001895.4) | 5 weeks / 1-3 years | Charles River (UK) | Commercial | 6 | All M | 10 weeks |
| Rabbit | *Oryctolagus cuniculus* | OryCun2.0 (GCA_000003625.1) | 5-6 months / 8-12 years | Envigo (UK) | Commercial | 6 | All M | 5 - 12 months |
| Pig (Domestic) | *Sus scrofa* | Sscrofa11.1 (GCA_000003025.6) | 6 months / 10-15 years | Harlan Ltd (UK) | Commercial | 3 | All M | 2 years |
| Dog (Beagle) | *Canis familiaris* | CanFam3.1 (GCA_000002285.2) | 1 year / 12-15 years | Harlan Ltd (UK) | Commercial | 4 | All M | 1 - 2.5 years |
| Cat | *Felis catus* | Felis_catus_9.0 (GCA_000181335.4) | 5-10 months / 15 years | Isoquimen Ltd (Spain) | Commercial | 6 | 2 F<br>4 M | 1 - 2.5 years |
| Horse (Welsh Mountain Pony) | *Equus caballus* | EquCab3.0 (GCA_002863925.1) | 12-15 months / 25-30 years | Animal Health Trust (UK) | Research colony | 4 | All M | 2 - 2.5 years |
| Grey Short-tailed Opossum | *Monodelphis domestica* | ASM229v1 (GCA_000002295.1) | 4-5 months / 4-8 years | Francis Crick Institute (UK) | Research colony | 5 | All M | 1 year |

**Table S2: All tissues used in the study**. Detailed description of all tissues samples used in the study, including the unique identifiers for three histone ChIP-seq libraries, the corresponding input libraries and RNA-seq libraries sequenced from each individual.

**Table S3: ChIP-seq mapping statistics**. The number of reads sequenced, mapped, passing quality control and after duplicate removal for all ChIP-seq and input libraries used in this study.

**Table S4: RNA-seq mapping statistics**. The number of sequenced, mapped, and multimapping RNA-seq reads for all libraries used in this study.

## Experimental model and subject details

The ten species used in this study were rhesus macaque (*Macaca mulatta*), common marmoset (*Callithrix jacchus*), mouse (C57BL/6J, *Mus musculus*), rat (Brown Norway, *Rattus norvegicus*), rabbit (*Oryctolagus cuniculus*), cat (*Felis catus*), dog (Beagle, *Canis familiaris*), horse (Welsh Mountain Pony, *Equus ferus*), pig (domestic pig, *Sus scrofa*), and grey short-tailed opossum (*Monodelphis domestica)*. All individuals used in this study were adults with no known health issues. Wherever possible, tissues from young adult males were used, however, some tissues were from females or older individuals. An overview of the origin, sex, and age for each animal used in the study is given in Table S1. The details for each individual animal and tissue used in this study are given in Table S2.

The use of all animals in this study was approved by the Animal Welfare and Ethics Review Board, under reference number NRWF-DO-02vs, and followed the Cancer Research UK Cambridge Institute guidelines for the use of animals in experimental studies. Tissues from seven species (macaque, marmoset, rabbit, cat, dog, horse, and opossum) were excess from routine euthanasia procedures (e.g., from individuals sacrificed during maintenance of research or breeding colonies.) Tissues from three species (mouse, rat, and pig) were purchased commercially (e.g., from animal research supply companies.)

### Macaque

Tissues from five rhesus macaques (*Macaca mulatta*) were used in this study. All individuals were male and ranged in age from 5 to 21 years (Table S2).

### Marmoset

Tissues from seven common marmosets (*Callithrix jacchus*) were used in this study. Five individuals were male and two were female. Individuals ranged in age from 1.5 to 19 years.

### Mouse

Tissues from twelve mice (C57BL/6J, *Mus musculus*) were used in this study. Eleven were 9-week-old males, and one individual was a 14-week-old female.

### Rat

Tissues from six rats (Brown Norway, *Rattus norvegicus*) were used in this study. All individuals were 10-week-old males.

### Rabbit

Tissues from six rabbits (*Oryctolagus cuniculus*) were used in this study. All individuals were male and ranged in age from 5 months to 1 year.

### Cat

Tissues from six cats (*Felis catus*) were used in this study. Four individuals were male and two were female. Individuals ranged in age from 1 to 2.5 years.

### Dog

Tissues from four dogs (Beagle, *Canis familiaris*) were used in this study. All individuals were male and ranged in age from 1 to 2.5 years.

### Horse

Tissues from four horses (Welsh Mountain Pony, *Equus ferus*) were used in this study. All individuals were male and ranged in age from 2 to 2.5 years.

### Pig

Tissues from three pigs (domestic pig, *Sus scrofa*) were used in this study. All individuals were 2-year-old males.

### Opossum

Tissues from six grey short-tailed opossums (*Monodelphis domestica)* were used in this study. All individuals were 1-year-old males.

## Method details

### Source and details of tissues

We performed ChIP-seq and RNA-seq on primary tissues isolated from 10 mammalian species. Primary tissues used were liver, skeletal muscle (from upper hind leg), brain (whole), and testis. Brain samples were representative of the whole brain (see details below) for most animals, with the exception of macaque, in which some of the brain regions were not available for use in this study (see Table S2). At least three independent biological replicates from different animals were used, with the only exception being H3K4me3 from horse testis, in which two of the replicates were from the same individual (Table S2). In most cases, matched tissues from the same individuals were used for all of the three ChIP-seq targets and RNA-seq (Table S2).

Tissues were prepared immediately post-mortem (typically within an hour) to maximize experimental quality. Tissues were processed by extracting the organ, dicing the tissue to small pieces and mixing it to get a fairly homogeneous mixture to give typical representation of the whole tissue (particularly important for whole brain samples). Tissues were then either immediately snap frozen on dry ice or liquid nitrogen (for RNA-seq), or formaldehyde crosslinked (see below) and then frozen on dry ice (for ChIP-seq).

### Chromatin immunoprecipitation and high-throughput sequencing (ChIP-seq)

Fresh, diced tissues were cross-linked in 1% formaldehyde in solution A (50 mM Hepes-KOH pH 7.5, 100 mM NaCl, 1 mM EDTA, 0.5 mM EGTA) for 20 minutes at room temperature, followed by addition of 2.5 M glycine solution to a final concentration of approximately 250 mM glycine and incubated for a further 10 minutes at room temperature to neutralize the formaldehyde. Samples were washed with cold PBS then frozen on dry ice and stored at −80°C until use.

Tissues were homogenized by either dounce homogenization of thawed tissues in PBS (for softer tissues from smaller species), or by grinding frozen tissues with a Qiagen TissueLyser II and stainless steel grinding jars, keeping the samples frozen by cooling jars in liquid nitrogen (for tissues from larger species or for muscle). After homogenization, samples were stored at −80 °C until further use.

Chromatin immunoprecipitations were done in Nunc deepwell (1 ml) 96-well plates. Each plate was set up to contain chromatin from 24 different tissue samples – each split into three different ChIP reactions (H3K4me3, H3K4me1 and H3K27ac) plus input – for a total of 96 samples (72 ChIP reactions plus 24 inputs) per plate. (As a result, all three ChIPs from the same tissue sample used the same input, except in cases where one of the ChIPs failed and needed to be repeated, in which case a new input was used for the new chromatin prep.) Tissue samples were assigned to 96-well plates semi-randomly, while maintaining a fairly even representation of species and tissue-type per plate. Sample position on the plates were also distributed in a semi-random fashion, while maximizing the distribution of samples with respect to species and tissue-type across the plate.

Antibodies were pre-bound to Protein G Dynabeads (Invitrogen). Antibodies used were H3K4me3 (Millipore 05-1339), H3K27ac (Abcam ab4729), and H3K4me1 (Abcam ab8895). Briefly, for each sample, 5 μg antibodies were pre-bound to 25 μl Protein G Dynabeads (Schmidt et al., 2009). Sufficient Dynabeads and antibodies (of the same type) were pooled for all 24 tissue samples, and incubated in 10 mL of block solution (1.5% BSA w/v in PBS) for at least 6 hours at 4 °C. Immediately prior to setting up the ChIP reactions, after chromatin extracts were prepared (see below), the antibody-bound beads were washed with 3x 10 mL block solution using a magnetic stand. Antibody-bound beads were then resuspended in block solution (enough for 100 μl per sample) and kept on ice.

Homogenized samples (24 at a time for a full 96-well plate) were lysed according to published protocols (Schmidt et al., 2009) to solubilize DNA-protein complexes. Approximately 0.3 to 0.5 g of homogenized tissue was lysed and resuspended in a final volume of 3 ml prior to sonication. Briefly, homogenized tissue was resuspended in 10 ml of lysis buffer 1 (50 mM Hepes-KOH pH 7.5, 140 mM NaCl, 1 mM EDTA, 10% glycerol, 0.5% NP-40, 0.25% Triton X-100) and incubated with rotation for 10 minutes at 4 °C. Samples were spun down at 2,500g for 3 minutes at 4 °C, and supernatants were decanted and discarded. The pelleted tissue was then resuspended in 10 ml of lysis buffer 2 (10 mM Tris-HCl pH 8.0, 200 mM NaCl, 1 mM EDTA, 0.5 mM EGTA) and incubated with rotation for 5 minutes at 4 °C. Samples were spun down at 2,500g for 3 minutes at 4 °C, and supernatants were decanted and discarded. Pelleted tissue was then resuspended in 3 ml lysis buffer 3 (10 mM Tris-HCl pH 8.0, 100 mM NaCl, 1 mM EDTA, 0.5 mM EGTA, 0.1% Na-Deoxycholate, 0.5% N-laurolsarcosine), transferred to a 5-ml Eppendorf tube, and incubated for at least 5 minutes (or up to 1 hour) prior to sonication. Protease inhibitors (Complete, EDTA-free, Roche, #11873580001) were added to all lysis buffers immediately prior to use.

Chromatin was fragmented to 300 bp average size by sonication on a Qsonica Q500 Sonicator with a 1/16” microtip at 40% amplitude for a total sonication time of 6 minutes (12 cycles of 30 seconds on, 60 seconds off). After sonication, 10% Triton X-100 was added to each sample to bring the total concentration of Triton X-100 to 1%. Samples were spun at 16,000g for 10 minutes at 4 °C, and the pellet was discarded to remove insoluble particles.

Each chromatin extract was split into three ChIP reactions: H3K4me3, H3K27ac, and H3K4me1. A small amount of extract (>3 μl) was reserved and stored at 4 °C to be used for the input (see below). Chromatin (800 μl per well, corresponding to approximately 0.1 g of homogenized tissue) and antibody-bound-beads (100 μl of suspension, equivalent to 5 ug of antibody, per well) were loaded into a 96-well Nunc deepwell 1 mL plate, and incubated overnight at 4 °C with end-over-end rotation.

Washes and subsequent steps were carried out with an Agilent Bravo liquid handling robot according to published protocols (Aldridge et al., 2013). Briefly, supernatant was discarded, and magnetic beads were washed 10x with 180 μl cold RIPA solution (50 mM Hepes-KOH pH 7.6, 500 mM LiCl, 1 mM EDTA, 1% NP-40, 0.7% Na-Deoxycholate), and then 2x with TBS. Magnetic beads were resuspended in 50 μl of elution buffer (50 mM Tris-HCl pH 8.0, 10 mM EDTA, 2% SDS), and incubated at 55 °C for 5 hours in a thermocycler to reverse crosslinks and elute from beads. Supernatants were removed from beads, diluted with equal volumes of TE buffer, and treated with RNase A (1 μl, Ambion #2271), followed by Proteinase K (1 μl, Invitrogen). 3 μl of chromatin extract (pre-ChIP) was added to elution buffer for the input samples and was reversed crosslinked, RNase and Proteinase K treated, and purified alongside the ChIP samples. Ampure bead purification was performed on the robot with a 1:1.8 DNA to Ampure bead ratio, and DNA was eluted from Ampure beads in 20 μl elution buffer. DNA concentration was measured with the Quant-iT dsDNA high-sensitivity kit on the PHERAstar microplate reader and was subsequently diluted to a concentration of 1 ng/μl. Illumina sequencing libraries were prepared from ChIP-enriched DNA or input DNA, using the ThruPLEX kit with 96 dual index adapters (Rubicon Genomics R400407) on the liquid-handling robot. Sequencing libraries were generally prepared from 10 ng (10 μl) of DNA, however amount of DNA ranged from 0.5 to 15 ng. Likewise, most libraries were amplified with 7 or 8 PCR cycles, but those with lower inputs of DNA into the library preparation were amplified with up to 16 PCR cycles. Libraries were run on an Agilent Tapestation 4200 with D1000 tapes for quantification. Libraries from each 96-well plate were mixed in equimolar concentrations into a single pool and sequenced on the Illumina HiSeq4000 with single end 50 bp reads.

### Total RNA sequencing (RNA-seq)

Total RNA was extracted from approximately 25 mg of snap-frozen tissue per sample. Tissue was thawed into 700 μl TRIzol and homogenized using a Precellys 24 tissue homogenizer with cooling system and 2 ml grinding tubes with beads (soft-tissue kit CK14 for liver, brain and testis, or the hard-tissue grinding kit MK28-R for muscle) for two intervals of 30 seconds. RNA was purified with phenol:chloroform extraction followed by isopropanol precipitation. RNA concentration was measured on the NanoDrop, samples were diluted, and 1-10 μg of RNA was taken forward in the procedure. RNA was treated with the TURBO DNA-free kit (Invitrogen) to remove any residual DNA. Illumina sequencing libraries were prepared using the Illumina TruSeq Stranded Total RNA with Ribo Gold kit (20020598) with Illumina RNA UD Indexes (20020492) according to the manufacturer’s protocol. Samples were run on Agilent Tapestation D1000 tapes to quantify sequencing libraries. Up to 12 libraries were combined into a single pool and sequenced on the Illumina NovaSeq 6000 for 300 cycles of paired end 150 bp reads.

## Quantification and statistical analysis

### Genome resources

The genome versions used in this study can be found in **Table S1: Species used in this study**. Briefly, all genomes were downloaded from the Ensembl version 98 (Yates et al., 2020) ftp as unmasked genomic DNA sequences, to facilitate the discovery of repetitive and transposable elements. For the mouse we used the primary assembly file, which excludes haplotypes and patches. All other species had no haplotype or patches, so we used the top-level DNA files.

### ChIP-seq mapping (Figures S1 and S2)

Reads were mapped to each species’ genome with BWA-MEM version 0.7.12 (Li and Durbin, 2009) using the default parameters. Low quality mapping reads were filtered out using SAMtools view version 1.3 with the -q1 flag (Li et al., 2009). Duplicates were removed with the Picard Tools MarkDuplicates program version 2.8.3 (https://broadinstitute.github.io/picard/). Mapping statistics were calculated using SAMtools flagstat version 1.3. To estimate the signal-to-noise ratio, we checked that the relative strand correlation (RSC) was above 0.8 for all libraries using Phantompeakqual tools version 1.14 (Landt et al., 2012). The mapping and RSC results are available in **Table S3: ChIP-seq mapping statistics**.

### ChIP-seq peak calling and signal saturation (Figures S1 and S2A)

To ensure that we have saturated the ChIP-seq signal for all libraries, we performed signal saturation tests (**Figure S2A**). With SAMtools view version 1.3, we subsampled quality filtered and duplicate removed reads from each biological replicate starting from 5 million reads to the maximum library depth, or to a maximum of 60 million reads, with a step of 5 million. For each subsampled set, we called enriched ChIP-seq regions using MACS2 version 2.1.1 (Zhang et al., 2008) using the broad peak mode (options: −q 0.05 --broad --broad-cutoff 0.1). An input library from the same individual and tissue (**Table S3: ChIP-seq mapping statistics**) and subsampled to the same sequencing depth was also used with MACS2. To discover biologically reproducible peaks, we looked for ChIP-seq peaks within replicates that overlapped with 50% of their length at least 50% of the peak of another replicate. Any such reproducible peaks appearing in at least two biological replicates were merged to produce the biologically reproducible set of histone enrichment peaks, while those not overlapping another replicate were not used for further analyses. Biologically reproducible H3K4me3 and H3K27ac reached ChIP-seq saturation at 20 million reads, while H3K4me1 reached saturation at 40 million reads (**Figure S2A**).

We used the ChIP-seq libraries for H3K27ac and H3K4me3 subsampled to 20 million reads for all further analyses. 12 of the somatic H3K4me3 libraries and one testis H3K4me3 library had less than 20 million reads after quality control and duplicate removal (**Table S3**), so we used all the reads from these libraries instead of subsamples. This did not reduce the total H3K4me3 peak numbers because H3K4me3 saturates at a sequencing depth below 20 million reads, especially in the somatic tissues (**Figure S2**). We subsampled all the H3K4me1 and matched input libraries to 40 million reads. The matched input sample for the macaque muscle library (unique identifier do17779) had around 21 million reads, which were used in MACS2 with the H3K4me1 library do17771.

### Capturing the signal across brain regions

To ensure that we are capturing the full complexity of the regulatory landscape in the brain, we compared our macaque H3K27ac ChIP-seq to a published study across three brain regions: cerebellum, cortex and subcortical structures (Vermunt et al., 2016). Across these three brain regions, they identified a total of 61,795 H3K27ac peaks in the macaque genome version rheMac3 while we found 85,025 H3K27ac peaks in whole macaque brain using genome version Mmul_10, suggesting that we effectively captured the brain regulatory landscape.

### Definitions of regulatory regions (Figure 1C)

Within each species and tissue, we defined regulatory regions from the overlap of biologically reproducible peaks (ChIP-seq peak calling and signal saturation (Figures S1 and S2A)). H3K27ac enrichment is characteristic of active regulatory elements (Heintzman et al., 2009; Heintzman et al., 2007; Rada-Iglesias et al., 2011). Concurrent H3K4me3 enrichment (Bernstein et al., 2005; Kim et al., 2005; Santos-Rosa et al., 2002; Schneider et al., 2004) in active regulatory region is characteristic of promoters, while concurrent H3K4me1 enrichment (Creyghton et al., 2010; Heintzman et al., 2009; Wang et al., 2008) is characteristic of enhancers. Therefore, we defined **active promoters** as H3K4me3 enriched regions that overlapped a H3K27ac enriched region with at least 50% of their length, regardless of whether H3K4me1 enrichment is also present. We took the length of the H3K4me3 peaks as the final active promoter region, but excluded the entire joint length of H3K27ac and H3K4me3 from further regulatory region calls. We identified as **active enhancers** H3K27ac histone enriched regions that overlap a H3K4me1 region with at least 50% of their lengths, keep only the span of H3K27ac peaks as the final active enhancer region. We excluded the whole region marked with H3K27ac and H3K4me1 from further regulatory region calls. Lastly, we defined H3K4me1 enriched regions that have no overlap with H3K27ac or H3K4me3 enriched regions as **primed enhancers**.

### Reannotation of genomes (Figure 1D)

For mouse and rat, we downloaded the available gene annotations from Ensembl version 98 (Yates et al., 2020). For all other species (macaque, marmoset, rabbit, pig, cat, dog, horse and opossum) we used a combination of our own RNA-seq data (Total RNA sequencing (RNA-seq)) and publicly available data to reannotate the genomes.

#### Transcript model generation

We generated gene annotations for each genome assembly using the previously described Ensembl annotation system (Aken et al., 2016). Briefly, we generated transcript models from multiple evidence sources taken from the public archives, using a variety of approaches: 1) mapping publicly available short read RNA-seq data from various tissues (search parameters: paired-end, >=75bp reads), including data generated by this study, 2) alignment of species-specific cDNAs (source: ENA (www.ebi.ac.uk/ena), obtained March 2019) to the genome and 3) protein-to-genome alignments of vertebrate UniProt (UniProt Consortium 2018) proteins with experimental evidence at the protein and transcript levels. In addition, whole genome alignments against the human GRCh38.p13 genome assembly were generated using LastZ (Harris, 2007) to identify regions of conserved synteny that subsequently guided mapping of conserved CDS exons from the GENCODE human gene set (Frankish et al., 2019). For pig and macaque, we obtained and mapped publicly available long read transcriptome data (PRJNA351265 and PRJNA320013 respectively) from various tissues to the genome using Minimap2 (Li, 2018).

#### Transcript filtering and prioritisation

For each locus, low quality transcript models with suboptimal mapping, limited intron-defining short read support or non-canonical splice sites were removed before collapsing and clustering non-redundant transcripts into gene models. We prioritized models generated from transcriptome data, having strong intron supporting evidence and high sequence identity (>90% coverage) to known vertebrate proteins. Gap filling was performed using homology data from projections to human annotations and mappings to UniProt proteins. To distinguish putative transcript isoforms from fragments, we assessed the coverage of protein alignments to each transcript relative to the size of the longest predicted open reading frame. Transcriptome data and cDNA alignments were used to extend models generated using homology data to annotate untranslated regions (UTR).

#### Gene model classification

We classified gene models into 3 main types: protein-coding, pseudogene and long non-coding RNA (lncRNA) using alignment qualities of all supporting data for each model. Models with alignments to known proteins, having little or no overlaps with repeat regions of the genome, having high intron support and well characterized canonical splice junctions were classified as protein coding. Pseudogenes were annotated by identifying genes with alignments to known proteins but with evidence of frame-shifting or located in repeat regions of the genome. Single-exon models with a corresponding multi-exon copy elsewhere in the genome were classified as processed pseudogenes. Gene models generated using transcriptomic data (short and long reads), lacking any protein supporting evidence and did not overlap a protein coding locus were classified as lncRNA.

Small non-coding RNA identification: Small non-coding (sncRNA) genes were added using annotations taken from RFAM (Griffiths-Jones et al., 2003) and miRbase (Griffiths-Jones et al., 2006). BLAST (Altschul et al., 1990) was run for these sequences to identify homologs in the genome sequence and models were evaluated for expected stem-loop structures using RNAfold (Lorenz et al., 2011). Additional machine learning based filters were applied to exclude predictions with sub-optimal alignments to the genome and non-conforming secondary structures. For other sncRNAs, models were built using the Infernal software suite (Nawrocki and Eddy, 2013).

### RNA-seq mapping and normalisation (Figure 1D and Table S4)

The RNA-seq reads were trimmed from adapters and for low quality bases using Trimmomatic version 0.33 (Bolger et al., 2014). For trimming, the TrueSeq3 paired end adapter sequences included with the Trimmomatic program were used. To remove low quality sequences from the reads, we removed those bases that had an average quality lower than 15 in a sliding window of four bases and the first and/or last three bases if below that threshold (options LEADING:3 TRAILING:3 SLIDINGWINDOW:4:15 MINLEN:36). For further analyses, we only kept reads with a minimum length of 36 bases, and only those that retained their paired read after trimming.

We mapped the trimmed RNA-seq reads using STAR version 2.6.0a (Dobin et al., 2013), the Ensembl 98 version of the genomes (Genome resources, **Table S1**), and gene annotation builds (Reannotation of genomes (Figure 1D)) to map each replicate RNA-seq library to known genes and transcripts. For STAR mapping, we used the following options: --outFilterType BySJout --outFilterMultimapNmax 100 --winAnchorMultimapNmax 100 --alignSJoverhangMin 8 --alignSJDBoverhangMin 1 --outFilterMismatchNmax 999 --outFilterMismatchNoverReadLmax 0.04 --alignIntronMin 20 --alignIntronMax 1000000 --quantMode GeneCounts --outSAMtype BAM SortedByCoordinate --outSAMstrandField intronMotif

To normalize the RNA-seq mapped libraries across replicates and tissues of the same species, we used Cufflinks version 2.2.1 (Trapnell et al., 2010). We first used the Cuffquant command specifying the strandedness of the library (option --library-type=fr-firststrand), followed by the Cuffnorm program treating each tissue as a sample, and each biological replicate as a replicate for that tissue. This produced normalized expression values for each annotated gene and transcript within a species and across all tissues. In **Figure 1D**, a gene or transcript was considered expressed in a tissue if this normalized value was above 0 FPKM.

### Validation of called regulatory regions (Figures S2B, S2C and S2D)

Mouse regulatory regions identified in the current study were compared to mouse regulatory regions annotated in Ensembl version 98 (Zerbino et al., 2016) and NCBI RefSeq functional elements (O’Leary et al., 2016) (downloaded 2017-09-26). We asked how many of the active promoters identified in the current study were annotated as promoters in either the Ensembl or RefSeq database. Given that neither external databases differentiate between enhancer types (i.e. active and primed enhancers) in an analogous manner to us, we combined primed and active enhancers identified in the current study into a single set. We than overlapped these enhancers with enhancers identified in either the Ensembl or RefSeq database. Overlap of any length in the genomic coordinates between a regulatory region identified in the current study and one annotated in the other database (Ensembl or RefSeq) was interpreted to mean the regulatory regions were common between the two sets, and lack of overlap was interpreted as a regulatory region specific to either the current study or to the other database (Ensembl or RefSeq). For simplicity, only regulatory regions mapping to chromosomes were considered for this analysis (those mapping to scaffolds were not considered). The resulting analyses are shown in **Figure S2B.**

For histone enrichment plots, local installations of deepTools version 3.3.1 (Ramirez et al., 2016) and WiggleTools (Zerbino et al., 2014) were used as follows: deepTools bamCompare was first used to subtract the corresponding input libraries from all quality controlled and duplicate removed (but not subsampled) ChIP-seq libraries and then WiggleTools mean to calculate average ChIP-seq enrichment across all mouse tissues and biological replicates for each histone mark. To create the heatmaps in **Figure S2D**, the resulting averages of read enrichment from H3K4me3, H3K27ac and H3K4me1 ChIP-seq libraries were compared with Ensembl Validated (i.e. overlap of our regions and Ensembl regulatory regions) and our novel regulatory mouse regions using the deepTools computeMatrix program with the options scale-regions --beforeRegionStartLength 2000 --afterRegionStartLength 2000 -missingDataAsZero --regionBodyLength 2000 --skipZeros and then plotted with the deepTools plotHeatmap program.

### Generating genome browser tracks (Figures 1A, 1B and 2A)

A biological replicate from a single individual for each species and tissue was arbitrarily chosen to display in the genome browser. Files were visualized in the IGV genome browser (Thorvaldsdóttir et al., 2012) with the appropriate genome and gene annotations files for each species. The genomic region around the genes encoding myosin heavy chains 1 and 2 (*Myh1* and *Myh2*) were extracted for each species and tissue from either bedGraph files (for ChIP-seq data), which represent read pileups for that biological replicate as generated by MACS2 (ChIP-seq peak calling and signal saturation (Figures S1 and S2A)) or wig files of uniquely mapping reads from the STAR alignments for the RNA-seq data (RNA-seq mapping and normalisation (Figure 1D and Table S4)). Bedgraph files were converted to the TDF file format with IGV tools to aid visualization in the browser. RNA-seq data are stranded, however the signal from the coding strand greatly dominated over the non-coding strand, and therefore only the coding strand was shown. Muscle samples visualized in **Figure 1A** were do17377 (macaque H3K4me3), do17393 (macaque H3K27ac), do18035 (macaque H3K4me1), do22674 (macaque RNA), do17664 (marmoset H3K4me3), do17690 (marmoset H3K27ac), do17715 (marmoset H3K4me1), do22678 (marmoset RNA), do15511 (mouse H3K4me3), do15539 (mouse H3K27ac), do15528 (mouse H3K4me1), do22610 (mouse RNA), do15941 (rat H3K4me3), do15952 (rat H3K27ac), do15918 (rat H3K4me1), do22601 (rat RNA), do17178 (rabbit H3K4me3), do17199 (rabbit H3K27ac), do17112 (rabbit H3K4me1), do22688 (rabbit RNA), do17356 (cat H3K4me3), do17365 (cat H3K27ac), do18036 (cat H3K4me1), do22638 (cat RNA), do17647 (dog H3K4me3), do17694 (dog H3K27ac), do17725 (dog H3K4me1), do22643 (dog RNA), do15887 (horse H3K4me3), do15926 (horse H3K27ac), do15954 (horse H3K4me1), do22676 (horse RNA), do17342 (pig H3K4me3), do16006 (pig H3K27ac), do16028 (pig H3K4me1), do26160 (pig RNA), do14518 (opossum H3K4me3), do14483 (opossum H3K27ac), do14565 (opossum H3K4me1), and do22663 (opossum RNA) (**Table S3** and **S4**). For mouse and dog, the same muscle samples from the same individuals were visualized in **Figure 1B**. Brain, liver and testis samples visualized in **Figure 1B** were do17085 (mouse brain H3K4me3), do17013 (mouse brain H3K27ac), do17044 (mouse brain H3K4me1), do22662 (mouse brain RNA), do15990 (mouse liver H3K4me3), do16031 (mouse liver H3K27ac), do16016 (mouse liver H3K4me1), do26179 (mouse liver RNA), do17048 (mouse testis H3K4me3), do17010 (mouse testis H3K27ac), do17079 (mouse testis H3K4me1), do22603 (mouse testis RNA), do17046 (dog brain H3K4me3), do17056 (dog brain H3K27ac), do17100 (dog brain H3K4me1), do22652 (dog brain RNA), do17397 (dog liver H3K4me3), do17324 (dog liver H3K27ac), do17341 (dog liver H3K4me1), do22650 (dog liver RNA), do17392 (dog testis H3K4me3), do17345 (dog testis H3K27ac), do17327 (dog testis H3K4me1), and do26151 (dog testis RNA).

### Intra-species cross-tissue activity (Figures 2B, S3A and S3B)

Within each species, we defined the tissue-specificity of regulatory regions by comparing the regulatory calls made within each of the tissues separately (Definitions of regulatory regions (Figure 1C)). Two regulatory regions were considered active across tissues if either overlapped another regulatory region of the same regulatory activity with at least 50% of its length. I.e. a tissue-shared active enhancer was considered tissue-shared only if it overlapped an active enhancer in another tissue (**Figure S3A**). All other combinations were considered intra-species dynamic (Intra-species dynamic regulatory signatures (Figure 4A)), and not included in any analyses or figures unless explicitly stated. A gene or transcript was considered expressed in a tissue according to the method outlined in RNA-seq mapping and normalization (Figure 1D and Table S4).

**Figure 2B** shows the sum across all ten species for the intersections between tissue activity using an UpSetR plot version 1.4.0 (Conway et al., 2017), while **Figure S3B** shows the data only for mouse. For all further analyses regulatory regions and gene expression were considered **tissue-specific** if they were only active in a single tissue, and **tissue-shared** if active in two or more tissues (**Figure S3A**).

### Association of enhancers to promoters (Figure 2C)

We used a distance rule to associate enhancers to promoters they might regulate, given that around 70% of enhancers do act on their nearest gene (Hait et al., 2018; Mifsud et al., 2015). Within each tissue, we associated primed and active enhancers called in that tissue to the nearest active promoter also called in that tissue. If there was no active promoter within 1 Mb of the enhancer, they were not assigned to an active promoter. We next tagged each active promoter, active and primed enhancer as tissue-specific or tissue-shared using the same rules as above (Intra-species cross-tissue activity (Figure 2B, S3A and S3B)) and defined an extra category for promoters – those promoters active across all four tissues were defined as **4-tissue-shared**. In **Figure 2C**, we show the distribution of the numbers of active and primed enhancers associated to tissue-specific and tissue-shared active promoters in each tissue. Promoters with more than 20 associated enhancers of any type are excluded from the graph, and a regression line representing the ration of tissue-specific to tissue-shared enhancers is shown. The data is a summary across all ten study species.

### Associations of regulatory regions to genes and differential gene expression analysis (Figure 2D)

For each active promoter within each tissue, we found the closest TSS in the given species using the Ensembl gene annotation created for this project (Reannotation of genomes (Figure 1D)). The TSS was defined as the start, i.e. most downstream coordinate, of a gene and that gene associated to a promoter if not further away than 1 Mb. Next, we used the enhancer-promoter association from above (Association of enhancers to promoters (Figure 2C)) to extend each active promoter associated gene to apply to that promoters’ enhancers.

We performed differential gene expression analyses between all other tissues and liver in a pairwise manner using DESeq2 version 1.10.1 (Love et al., 2014) with default parameters and a Benjamini-Hochberg adjusted p-value threshold of 0.05 (padj < 0.050000). As input for DESeq2 we used raw read counts produced with the STAR aligner (RNA-seq mapping and normalisation (Figure 1D and Table S4)). Specifically, for each species, we tested the differential expression between muscle, brain or testis against the liver and in **Figure 2D** report log2fold changes that passed the thresholds. Within each tissue, we further created a more stringent subset from all tissue-shared regulatory regions (Intra-species cross-tissue activity (Figure 2B, S3A and S3B)) to only include those shared across all four tissues (**4-tissue-shared**).

### Whole genome alignments

For whole genome alignment between the eutherian mammals (macaque, marmoset, mouse, rat, rabbit, pig, cat, dog and horse) we used the EPO eutherian mammal alignments from Ensembl version 98 (Herrero et al., 2016). For whole genome alignments between the eutherian mammals and opossum, we used the PECAN alignments also from Ensembl version 98. We aligned all species to mouse in a pairwise manner, first from all other species to mouse and then from mouse to all other species. A regulatory region was considered **maintained** (*P_Mi_*; see equation 1 and **Figure S3A**) if it overlapped another regulatory region of any type with at least one base. A regulatory region was considered **recently evolved** (*P_Ri_*; see equation 2 and **Figure S3A**) if it was aligned to other species but did not overlap a regulatory region in any of them, or if it was not aligned to any other species. Any regulatory region aligned to multiple locations was excluded from further analyses, i.e. only 1-to-1 alignments were kept.

Equations demonstrating the computation of cross-species conservation of promoters are shown in the following sections. The same operations were computed for active and primed enhancers but are not shown here.

### The recently evolved and maintained regulomes across species (Figure 3A)

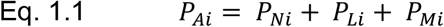

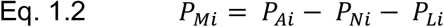

*P_Ai_* – number of all active promoters in species *i*

*P_Ni_* – number of active promoters in species *i* with no alignment to any other species

*P_Li_* – number of active promoters in species *i* with an alignment to any other species, but no regulatory region aligned in any other species

*P_Mi_* – number of active promoters in species *i* with an alignment of at least one base length to any regulatory region in any other species

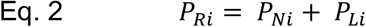

*P_Ri_* – number of recently evolved active promoters in species *i*

**Figure 3A** Shows the **∑ *P_Mi_*** (Eq. 1.2) and **∑ *P_Ri_*** (Eq. 2) across all ten species for active promoters, and analogous calculation for active and primed enhancers.

### Pairwise comparisons between species (Figures 3B and 3C)

We performed two pairwise comparisons, the first stratifying regulatory regions by tissue-shared and tissue-specific (**Figure 3B**), and the second further stratifying tissue-specific regulatory regions by their tissue of activity (i.e. liver-specific, muscle-specific, brain-specific and testis-specific) (**Figure 3C**). The stratification was limited to the identity of the query regulatory region, but the query region was considered maintained if it aligned to any regulatory region in the other species (regardless of tissue-specificity). For example, a liver-specific mouse active promoter was consider maintained and counted as tissue-specific (**Figure 3B**) and liver-specific (**Figure 3C**) in all the following cases: 1) if it aligned to a liver-specific active promoter, 2) if it aligned to a tissue-shared active promoter, 3) if it aligned to a muscle-specific active promoter, or 3) if it aligned to active or primed enhancers of any tissue-specificity in the other species. For definitions of tissue-specificity see Methods section Intra-species cross-tissue activity (Figure 2B, S3A and S3B).

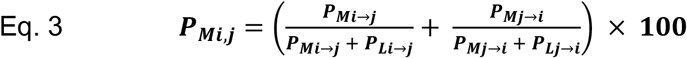

*P_Mi,j_* – fraction of alignable active promoters with activity in both species *i* and *j*, i.e. aligned to a regulatory active region. Defined as maintained regulatory regions. See also Eq. 6

*P_Mi→j_* – number of active promoters in species *i* with an alignment of at least one base length to any regulatory region in species *j*.

*P_Li→j_* – number of active promoters in species *i* with an alignment of at least one base length to species *j,* but not aligned to a regulatory region

To calculate the zero point, we generated a fourth ChIP-seq replicate for all histone modifications for mouse and cat (**Table S3**) and called peaks using the same methods as for other replicates (ChIP-seq peak calling and signal saturation (Figures S1 and S2A)). We then used all possible combinations of three replicates to estimate interindividual reproducibility (Definitions of regulatory regions (Figure 1C)) as a measure of both the variation between individuals and biases introduced by our analyses.

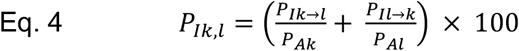

*P_IK,l_* – fraction of active promoters with regulatory activity between a pair of biological replicates *k* and *l*, i.e. reproducible between individuals.

*P_IK→l_* – number of active promoters in individual *k* that overlap a regulatory active region in individual *l* by at least one base

*P_Ak_* – total number of active promoters in individual *k*

**Figures 3B** and **3C** show *P_Mi,j_* (Eq. 3) for every pair of species at divergence > 0 MYA (45 comparisons) and *P_Ik,l_* (Eq. 4) at divergence = 0 for every pair of 4 mouse and every pair of 4 cat biological replicates (12 comparisons). For **Figure 3B** we first plotted regions identified as tissue-shared in species *i* and species *j*, and then regions identified as tissue-specific in species *i* and species *j.* Similarly, for **Figure 3C** we plotted separately all tissue-specific regions depending on which tissue they were active in. We plotted the resulting graphs in R version 3.6.2 (R Core Team, 2019) using ggplot2 version 3.1.1 (Wickham, 2016), and performed linear regression using the geom_smooth() ggplot2 method. To test for statistical significance, we fitted the data to a linear model using the R function lm() and tested the resulting linear models using the built-in anova() function for the interaction of divergence time and tissue-specificity. Specifically, for **Figure 3B** we report the two-way ANOVA p-value for the interaction of divergence time (factor 1) and regions identified as active promoters and active enhancers (factor 2); for the interaction of divergence time (factor 1) and regions identified as active promoters and primed enhancers (factor 2); and for the interaction of divergence time (factor 1) and regions identified as active enhancers and primed enhancers (factor 2). For **Figure 3C** we first report the two-way ANOVA p-value for the interaction of divergence time (factor 1) and active promoters identified as testis-specific or any other tissue-specific region (factor 2). We then report the two-way ANOVA p-value for the interaction of divergence time (factor 1) and active promoters being identified as tissue-specific or tissue-shared (factor 2). All divergence times between species were taken from Ensembl Compara version 98 (Herrero et al., 2016).

### Intra-species dynamic regulatory signatures (Figures 4A and S3A)

To define regulatory regions that change regulatory identity between the tissues of a species, we performed cross-tissue overlap as described for determining tissue-specific and tissue-shared regions (Intra-species cross-tissue activity (Figure 2B, S3A and S3B)). Briefly, if regulatory regions of different identities overlapped each other with at least 50% of their length between tissues, we called these regions **intra-species dynamic** (**Figure S3A**). Specifically, overlap of an active promoter in one tissue to either an active or primed enhancer in another tissue was called a **dynamic promoter/enhancer (dynamic P/E)**, while the overlap of an active enhancer in one tissue to a primed enhancer in another tissue was called a **dynamic enhancer (dynamic E)**. The sum of all dynamic regions, and non-dynamic regions, across all ten species is shown in **Figure 4A.**

### Evolutionary dynamic regulatory signatures (Figures 4B and S3A)

For all maintained regulatory regions (*P_Mi,j_* Eq. 3, The recently evolved and maintained regulomes across species (Figure 3A), **Figure S3A**), we next asked how the regulatory signature changes through evolution. For each pairwise comparison between species, we counted how many regulatory regions of one identity aligned with at least one base to a regulatory region of all other signatures. These calculations were limited only to maintained regulatory regions which only align to one other regulatory region between species. The text below shows the calculation for active promoters as an example, but all other regulatory regions were calculated similarly.

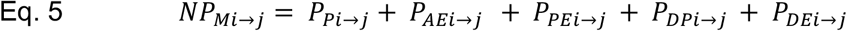

*NP_Mi→j_* – total number of maintained promoters in species *i* when compared to species *j*

*P_pi→j_* – Total number of active promoters in species *i* with a 1-to-1 alignment to at least one base of an active promoter in species *j;* these represent **evolutionarily stable promoter signatures**

*P_AEi→j_* – Total number of active promoters in species *i* with a 1-to-1 alignment to at least one base of an active enhancer in species *j;* these represent **evolutionarily dynamic promoter signatures**

*P_PEi→j_* – Total number of active promoters in species *i* with a 1-to-1 alignment to at least one base of a primed enhancer in species *j;* these represent **evolutionarily dynamic promoter signatures**

*P_DPi→j_* – Total number of active promoters in species *i* with a 1-to-1 alignment to at least one base of an intra-species dynamic promoter region in species *j;* see also Intra-species dynamic regulatory signatures (Figure 4A and S3A)

*P_DEi→j_* – Total number of active promoters in species *i* with a 1-to-1 alignment to at least one base of an intra-species dynamic enhancer region in species *j;* see also Intra-species dynamic regulatory signatures (Figure 4A and S3A)

**Figure 4B** Shows the ∑(*P_Pi→j_* + *P_AEi→j_* + *P_PEi→j_* + *P_DPi→j_* + *P_DEi→j_*) for all pairs of species for the active promoters, and analogous calculations for the other regulatory regions, in a Circos plot (Krzywinski et al., 2009).

### Evolutionary dynamic regulatory identities across divergence time (Figure 4C)

Next, we asked if evolutionary dynamics of promoter and enhancer signatures is correlated to the divergence time between species. For this, we focused on all maintained regulatory regions (The recently evolved and maintained regulomes across species (Figure 3A)) between pairs of species, and asked how often they align to a regulatory region with another regulatory signature (Evolutionary dynamic regulatory signatures (Figure 4B and S3A)). The comparisons were limited to 1-to-1 aligned regulatory regions.

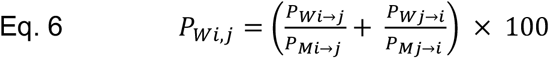

*P_Wi,j_* – fraction of maintained active promoters switching regulatory activity between species *i* and *j*, i.e. aligned to a regulatory active region. Defined as **evolutionarily dynamic promoter signatures.** See also Eq. 5

*P_Wi→j_* – number of active promoters in species *i* aligned to an active or primed enhancer in species *j*

*P_Wi→j_* – number of active promoters in species *i* aligned to any regulatory region in species *j;* see also Eq. 3

*P_Wi→j_* – number of active promoters in species *j* aligned to an active or primed enhancer in species *i*

*P_Wi→j_* – number of active promoters in species *j* aligned to any regulatory region in species *i;* see also Eq. 3

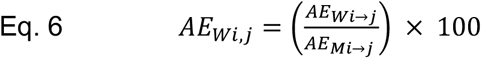

*AAE_Wi,j_* – fraction of maintained active enhancers switching regulatory activity between species *i* and *j*, i.e. aligned to a regulatory active region. Defined as **evolutionarily dynamic enhancer signatures.** See also Eq. 5

*AE_Wi→j_* – number of active enhancers in species *i* aligned to a primed enhancer in species *j*

*AE_Wi→j_* – number of active enhancers in species *i* aligned to any regulatory region in species *j;* see also Eq. 3

**Figure 4C** Shows the *P_W,j_* and *AE_Wi,j_* between all pairs of species (45 comparisons) for every pair of species at divergence > 0 MYA (45 comparisons) and at divergence = 0 the average intra-species dynamic activity (Intra-species dynamic regulatory signatures (Figure 4A and S3A)). All divergence times between species were taken from Ensembl version 98 (Herrero et al., 2016). We plotted the resulting graphs in R version 3.6.2 (R Core Team, 2019) using ggplot2 version 3.1.1 (Wickham, 2016), and performed linear regression using the geom_smooth() ggplot2 method.

### Outgroup analysis (Figures 4D, S4B and S4C)

Whole genome alignments of regulatory regions were parsed to get all 1-to-1 alignments for mouse/rat/rabbit and separately for cat/dog/horse (see Evolutionary dynamic regulatory signatures (Figure 4B and S3A)). Only genomic regions that were maintained as either an active promoter, active enhancer, or primed enhancer in all three of the species in the triad (mouse/rat/rabbit and cat/dog/horse) were considered in this analysis. Genomic regions identified as an intra-species dynamic region in any of the three species were excluded from this analysis for simplicity. Analyses were done separately for each triad. The overall proportion of active promoters, active enhancers, and primed enhancers in the outgroup species (rabbit or horse) for the genomic regions considered is shown as “All” in (**Figures 4D** and **S4**). Given the combination of regulatory signatures in the ingroup species (mouse/rat or cat/dog), we asked what the identify was in the outgroup species (rabbit or horse, respectively). For example, in the AP/AE situation, this could represent either an active promoter in mouse and an active enhancer, or vice versa. The regulatory signature of the genomic region in rabbit would then be queried. Percentages and raw numbers are shown separately for each triad in **Figure S4B** and **Figure S4C**. The combined numbers and percentages are also shown in the bottom panels of **Figure S4B** and **Figure S4C** and in **Figure 4D**. Chi-square two-tailed tests (degrees of freedom = 2) were used to test whether outgroup distributions of active promoters, active enhancers and primed enhancers for each ingroup combination differed statistically from the background (“All”) distribution (**Figure 4D**). Expected values were calculated based on the percentages of active promoters, active enhancers, and primed enhancers in the background.

### Model of evolutionary dynamics between regulatory regions (Figure 4E)

For the model of all possible evolutionary dynamics between active promoters (*AP*), active enhancers (*AE*) and primed enhancers (*PE*), we extracted probabilities using the observed frequencies in the triad analysis above (Outgroup analysis (Figure 4D, S4B and S4C)). For each regulatory region we calculated the relative probabilities of retaining the same signature (*AP* → *AP*, *AE* → *AE* and *PE* → *PE*), and changing to a regulatory region of another signature (for example: *AP* → *AE* or *AE* → *AP*). The probabilities were calculated from the observed frequencies in both outgroup analyses combined (mouse/rat/rabbit and cat/dog/horse), using only those evolutionary relationships where parsimony could be used to determine the ancestral state as a single regulatory region signature.

Specifically, for active promoters:

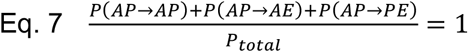

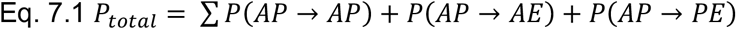

*P*(*AP* → *AP*) – probability of an ancestral active promoter remaining an active promoter. Observed frequency from triad relationship ingroups AP/AP and outgroup AP.

*P*(*AP* → *AE*) – probability of an ancestral active promoter evolving to an active enhancer. Observed frequency from triad relationship ingroups AP/AE and outgroup AP.

*P*(*AP* → *PE*) – probability of an ancestral active promoter evolving to a primed enhancer. Observed frequency from triad relationship ingroups AP/PE and outgroup AP.

Specifically, for active enhancers:

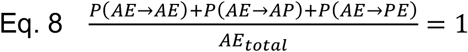

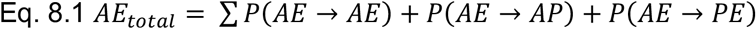

*P*(*AE* → *AE*) – probability of an ancestral active enhancer remaining an active enhancer. Observed frequency from triad relationship ingroups AE/AE and outgroup AE.

*P*(*AE* → *AP*) – probability of an ancestral active enhancer evolving to an active promoter. Observed frequency from triad relationship ingroups AE/AP and outgroup AE.

*P*(*AE* → *PE*) – probability of an ancestral active enhancer evolving to a primed enhancer. Observed frequency from triad relationship ingroups AE/PE and outgroup AE.

Specifically, for primed enhancers:

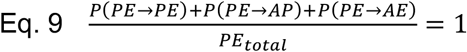

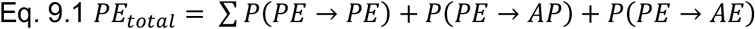

*P*(*PE* → *PE*) – probability of an ancestral primed enhancer remaining a primed enhancer. Observed frequency from triad relationship ingroups PE/PE and outgroup PE.

*P*(*PE* → *AP*) – probability of an ancestral primed enhancer evolving to an active promoter. Observed frequency from triad relationship ingroups PE/AP and outgroup PE.

*P*(*PE* → *AE*) – probability of an ancestral primed enhancer evolving to an active enhancer. Observed frequency from triad relationship ingroups PE/AE and outgroup PE.

**Figure 4E** shows the resulting calculations for all evolutionary relationships as numbers above the arrows.

### ChIP-seq and RNA-seq read enrichment around evolutionarily dynamic regulatory regions (Figures 4F and 4G)

To validate evolutionary switching from active enhancer to active promoter (evolutionary dynamic P/Es, **Figure S3A**), we selected a subset of regulatory regions from the outgroup analysis (Outgroup analysis (Figures 4D, S4B and S4C)) that are most likely to represent a true evolutionary switch. Specifically, we only chose those regions that had an active promoter signature in one ingroup, active enhancer signature in the other ingroup and an active enhancer signature in the outgroup as the most parsimonious conclusion is that the ancestral state was an active enhancer.

For **Figure 4F**, we separated these regions within each outgroup species into sets that showed active promoter signature (“AP Dynamic”) and active enhancer signature (“AE Dynamic”). We then selected from the same species the same number of control regions as those that never show evolutionarily dynamic activity. We next generated the per-species averages of ChIP-seq enrichment across these regions for all replicates of a species as described before (Validation of called regulatory regions (Figures S2B, S2C and S2D)) but extending the flanking regions to 10 Kb. Finally, we averaged across all per-species averages to generate the graphs in Figure 4F.

For **Figure 4G** we used the same AP Dynamic and AE Dynamic sets as described above, but only the active enhancers as control regions. For RNA-seq enrichment plots, we used local installations of deepTools version 3.3.1 (Ramirez et al., 2016) and WiggleTools mean (Zerbino et al., 2014) to calculate the maximum RNA-seq enrichment across all biological replicates across all tissues in a species. For **Figure 4G**, the resulting maximum RNA-seq values were compared between evolutionarily dynamic and control regions using the deepTools computeMatrix program with the options scale-regions --beforeRegionStartLength 10000 --afterRegionStartLength 10000 --missingDataAsZero --regionBodyLength 2000 -skipZeros within each species. To generate the resulting boxplots in Figure 4G, all species’ values were combined. The p-values were calculated with the ggpubr package in R (Kassambara, 2020), using the stat_compare_means function using a t-test testing if active promoters (AP) had higher expression than active enhancers (AE) than control regions.

### Tissue-specificity of evolutionarily dynamic regions (Figure 4H)

To create the tissue-specificity UpSetR version 1.4.0 plots (Conway et al., 2017) in **Figure 4H**, we extracted all regions corresponding to **evolutionary dynamic promoter signatures** (Figure S3A, Evolutionary dynamic regulatory signatures (Figures 4B and S3A)). Specifically, we extracted those active promoters that have a 1-to-1 alignment to an active or primed enhancer in any other species and active and primed enhancers that have a 1-to-1 alignment to an active promoter in another species. We then considered the tissue-specificity of the regions categorising them according to the regulatory identity in the species they were extracted from. For example, for a region identified as an active promoter in rat and an active enhancer in rabbit we considered its tissue-specificity only in rat for the active promoter category and tissue-specificity only in rabbit for the active enhancer category. We plotted the within species tissues specificity of those selected regulatory regions as outlined in Methods section Intra-species cross-tissue activity (Figures 2B, S3A and S3B). We performed all statistical tests using the binom.test() function in R version 3.6.2 (R Core Team, 2019).

### Repeat masking and classification of transposable elements

To identify transposable elements in all genomes we used RepeatMasker version open-4.0.7 (Smit, 2013-2015) using the crossmatch search engine and RepBase Release 20170127 (Bao et al., 2015). For each species we ran RepeatMasker in the default mode, specifying the species’ scientific name (Macaque, “macaca mulatta”; Marmoset, “callithrix jacchus”; Mouse, “mus musculus”; Rat, “rattus norvegicus”; Rabbit, “oryctolagus cuniculus”; Pig, “sus scrofa”; Dog, “canis familiaris”; Cat, “felis catus”; Horse, “equus caballus”; Opossum, “monodelphis domestica”). For figures, we used DNA, LINE, LTR and SINE categories from RepBase annotation. These correspond to different levels of transposable element classification hierarchies (Bourque et al., 2018), but still represent exclusive non-overlapping sets of the hierarchy. Namely, DNA corresponds to the DNA transposons class, while all other groups belong to the retrotransposon classes. The LTRs are a subclass of retrotransposons, while the LINEs and SINEs are superfamilies of the non-LTR subclass of retrotransposons.

### Relative transposable element enrichment of tissue-specific and tissue-shared recently evolved regulatory and maintained regions (Figures 5A and S5A)

We first extracted all **recently evolved** and **maintained regulatory regions** (**Figure S3A**, The recently evolved and maintained regulomes across species (Figure 3A)). Next, within each species we calculated the number of tissue-specific recently evolved regions that overlap with at least one base subgroups of transposable elements as defined by RepBase (Repeat masking and classification of transposable elements). For example, for tissue-specific active promoters overlapping any LINE with at least one base, we counted the number occurring in all possible subgroups (for example: L1, L2, CR1). We next repeated the same process for tissue-shared active promoters overlapping any LINE. To generate the relative enrichment shown in **Figure 5A** and **S5A**, we calculated the percent of tissue-specific and tissue-shared regulatory regions overlapping specific subgroups by dividing each subgroup count by the total counts for that group and multiplying by 100. For example, for L1 we divided the number of tissue-specific active promoters overlapping an L1 by the total number of tissue-specific active promoters overlapping any LINE. Similarly, for the tissue-shared we divided the number of tissue-shared active promoters overlapping an L1 by the total number of tissue-shared active promoters overlapping any LINE. Finally, we subtracted the tissue-shared portions with the tissue-specific to generate a relative enrichment. Consequently, positive values indicate a higher proportion of that subgroup in the tissue-specific than in the tissue-shared. For example, pig has a value of 16 for L1 active promoters because 47% of all tissue-specific active promoters overlapping a LINE belonged to the L1 subgroup, compared to 31% of the tissue-shared active promoters. The heatmap of all relative enrichments in **Figures 5A** and **S5A** were generated using the heatmap.2() function in gplots package version 3.0.1.1 (Warnes et al., 2019). The p-values were calculated on the original counts, using the Z-test in R base function prop.test and bonferonni correction using the total number of tests across the matrix implemented in R base function p.adjust. For generating the heatmaps images, we filtered all possible subgroups to include only those that have at least 100 tissue-specific and 100 tissue-shared occurrences in any species, and manually refined the selection to only include those that are informative for multiple linages, but total counts across all subgroups were used for the p-value calculations. **Figure 5A** shows the relative transposable element enrichments for **recently-evolved** regulatory regions, while **Figure S5A** shows the enrichments for **maintained regulatory regions**.

### ChIP-seq and RNA-seq read enrichment around LINE-associated recently evolved regulatory regions (Figures 5B and 5C)

To examine the raw signal surrounding LINE-associated active promoters, we first extracted recently evolved active promoters overlapping a LINE L1 or LINE L2 with at least one base (overlap defined in Relative transposable element enrichment of tissue-specific and tissue-shared recently evolved regulatory and maintained regions (Figure 5A and S5A)). We next chose those active promoters that had overlap with LINE L2s and have tissue-shared activity (as defined in Intra-species cross-tissue activity (Figures 2B, S3A and S3B)). For those active promoters that had overlap with LINE L1s and were tissue-specific, we further subdivided them by the tissue of activity. For **Figure 5B**, we generated the per-tissue averages of ChIP-seq enrichment across all replicates of a species as described before (Validation of called regulatory regions (Figures S2B, S2C and S2D)) but making an average across all replicates of a tissue within a species and extending the flanking regions to 10 Kb. Finally, we combined averaged across all per-species averages to generate the graphs in Figure 5B.

For RNA-seq enrichment plots, we used local installations of deepTools version 3.3.1 (Ramirez et al., 2016) and WiggleTools max (Zerbino et al., 2014) to calculate the maximum RNA-seq enrichment across all biological replicates of the tissue. For **Figure 5C**, the resulting maximums RNA-seq values were compared to LINE associated active promoters using the deepTools computeMatrix program with the options scale-regions --beforeRegionStartLength 10000 --afterRegionStartLength 10000 --missingDataAsZero --regionBodyLength 2000 – skipZeros within each species. To generate the resulting boxplots in Figure 5C, all species’ values were combined. The p-values were calculated with the ggpubr package in R (Kassambara, 2020), using the stat_compare_means function using a t-test testing if the mean of each boxplot is significantly different from all other expressed in that tissue (i.e. within rows).

### Age of LINEs (Figures 5D and S5B)

To estimate the age of LINEs, we used the percent mutations from RepBase consensus sequences of each element as reported by RepeatMasker (Repeat masking and classification of transposable elements). We extracted all LINE L2 and L1 matches in the genome and characterized them as **regulatorily inactive** if they did not overlap a regulatory region we identified in this project, as **recently evolved regulatory region** if they were recently evolved (The recently evolved and maintained regulomes across species (Figure 3A)), and **evolutionarily dynamic regulatory signature** if we had found them to align to a regulatory region of another signature (Evolutionary dynamic regulatory signatures (Figures 4B and S3A)). For **Figure 5D** we plotted the mutations for all species combined, while **Figure S5** shows the same data but split by the species it was identified in. The p-values were calculated with the ggpubr package in R (Kassambara, 2020), using the stat_compare_means function and a one sided Wilcoxon test between all pairs of categories (regulatorily inactive, recently-evolved and evolutionarily-dynamic regulatory region).

### Relative transposable element enrichment of evolutionarily dynamic and stable regulatory signatures (Figure 5E)

We first extracted all evolutionarily dynamic regulatory regions, i.e. evolutionarily dynamic promoters and evolutionarily dynamic enhancers (Evolutionary dynamic regulatory signatures (Figures 4B and S3A)). To calculate the relative enrichment of evolutionarily dynamic regions (**switch** regulatory regions) to those not found to be evolutionarily dynamic (**stable** regulatory regions), we performed calculations similar to the recently evolved relative enrichment (Relative transposable element enrichment of tissue-specific and tissue-shared recently evolved regulatory and maintained regions (Figure 5A and S5A)) but changing the groups of regulatory regions being compared. To generate the relative enrichment shown in **Figure 5E**, for each category of regulatory region we subtracted the percentage of stable regulatory regions belonging to a subgroup from the percentage of evolutionarily dynamic regulatory regions. Consequently, positive values indicate a higher proportion of that subgroup in the evolutionarily dynamic than in stable regulatory regions. For example, rat has a value of −11 for L1 active enhancers because 63% of all evolutionarily dynamic active enhancers overlapping a LINE belonged to the L1 subgroup, compared to 74% of the stable active enhancers. The heatmap of all relative enrichments in **Figure 5E** was generated using the heatmap.2() function in gplots package version 3.0.1.1 (Warnes et al., 2019). The p-values were calculated on the original counts, using the Z-test in R base function prop.test and Bonferroni correction using the total number of tests across the matrix implemented in R base function p.adjust. For generating the heatmaps images, we filtered all possible subgroups to include only those that have at least 100 tissue-specific and 100 tissue-shared occurrences in any species, and manually refined the selection to only include those that are informative for multiple linages, but total counts across all subgrops were used for the p-value calculations.

### Constrained element content in tissue-specific LINEs (Figures S5C and S5D)

To examine the difference in sequence constraint within regulatorily active LINE transposable elements and avoid bias, we focused on those LINE elements that overlapped tissue-specific regulatory regions in all species. For **Figure S5C** for each tissue-specific regulatory region associated with a LINE L1 or L2 we extracted the GERP rejected substation scores (Cooper et al., 2005) from Ensembl version 98 (Herrero et al., 2016). Briefly, unalignable genomic regions do not have a GERP score, while a negative score in alignable indicates more sequence constraint than expected and a positive score indicates less. We plotted the resulting graphs in R version 3.6.2 (R Core Team, 2019) using ggplot2 version 3.1.1 (Wickham, 2016), and calculated p-values using the base R wilcox.test.

For **Figure S5D** we extracted the number of constrained elements, i.e. short genomic regions that have more sequence constraint than expected (Cooper et al., 2005), for each tissue-specific regulatory region associated with a LINE L1 or L2 from Ensembl version 98 (Herrero et al., 2016). Unlike the analysis reported in **Figure 5C**, this includes also unaliganble genomic regions. We plotted the resulting graphs in R version 3.6.2 (R Core Team, 2019) using ggplot2 version 3.1.1 (Wickham, 2016), and calculated p-values using the base R chisq.test.

